# Task-specific neural processes underlying conflict resolution during cognitive control

**DOI:** 10.1101/2022.01.16.476535

**Authors:** Yuchen Xiao, Chien-Chen Chou, Garth Rees Cosgrove, Nathan E Crone, Scellig Stone, Joseph R Madsen, Ian Reucroft, Yen-Cheng Shih, Daniel Weisholtz, Hsiang-Yu Yu, William S. Anderson, Gabriel Kreiman

## Abstract

Cognitive control involves flexibly combining multiple sensory inputs with task-dependent goals during decision making. Several tasks have been proposed to examine cognitive control, including Stroop, Eriksen-Flanker, and the Multi-source interference task. Because these tasks have been studied independently, it remains unclear whether the neural signatures of cognitive control reflect abstract control mechanisms or specific combinations of sensory and behavioral aspects of each task. To address this question, here we recorded invasive neurophysiological signals from 16 subjects and directly compared the three tasks against each other. Neural activity patterns in the theta and high-gamma frequency bands differed between incongruent and congruent conditions, revealing strong modulation by conflicting task demands. These neural signals were specific to each task, generalizing within a task but not across tasks. These results highlight the complex interplay between sensory inputs, motor outputs, and task demands and argue against a universal and abstract representation of conflict.

## Introduction

The ability to flexibly route information is central to daily activities, especially when faced with a complex and conflicting interplay of sensory information, choices, and goals. Cognitive control refers to the ability to regulate actions toward achieving overriding goals. Cognitive control is mentally effortful because of the need to suppress autonomous responses toward salient but goal-irrelevant stimulus attributes (Gratton et al., 1992; Miller and Cohen, 2001). Such costs are unavoidable for successful adaptation to various environments (Diamond, 2013). Impairment in cognitive control is associated with a wide range of mental disorders, including addiction, depression, and schizophrenia (Goschke, 2014; Lesh et al., 2011; Zilverstand et al., 2018). An essential component of cognitive control is conflict resolution, which entails mental operations involving conflict detection and monitoring (Botvinick et al., 2001), response selection and inhibition (Goghari and MacDonald, 2009), performance monitoring and evaluation (Ridderinkhof et al., 2004), and error-detection (Fu et al., 2019; Ridderinkhof et al., 2004; Tang et al., 2016).

Many experimental tasks have been used to study cognitive control during conflict resolution. Paradigmatic examples include the Stroop task (Stroop, 1935), the Eriksen-Flanker task (referred to as “Flanker” throughout the text, (Eriksen and Eriksen, 1974), and the Multi-Source Interference task (MSIT, referred to as “Number” throughout the text, (Bush and Shin, 2006)). Common to all these tasks is the comparison between a congruent condition and an incongruent condition (**Figure 1**). In the Stroop task, subjects name the font color of a color word (e.g., “red,” “green,” “blue”) when the semantic meaning of the word agrees (congruent condition) or disagrees (incongruent condition) with its font color. In the Flanker task, subjects have to recognize a symbol such as a letter or an arrow, embedded among the same symbols (congruent condition) or different symbols (incongruent condition) (Davelaar and Stevens, 2009; Eriksen and Eriksen, 1974; Mayr et al., 2003). The multi-source interference task (Bush and Shin, 2006) combines multiple dimensions of cognitive interference from the Stroop, Flanker, and Simon (Simon and Berbaum, 1990) tasks. The MSIT stimulus consists of three numbers (chosen from 0, 1, 2, or 3) in which one number (target) is always different from the other two numbers (distractors). Subjects are instructed to identify the target number under conditions where it is congruent (e.g., 100) or incongruent (e.g., 313) with its position.

**Figure 1.**
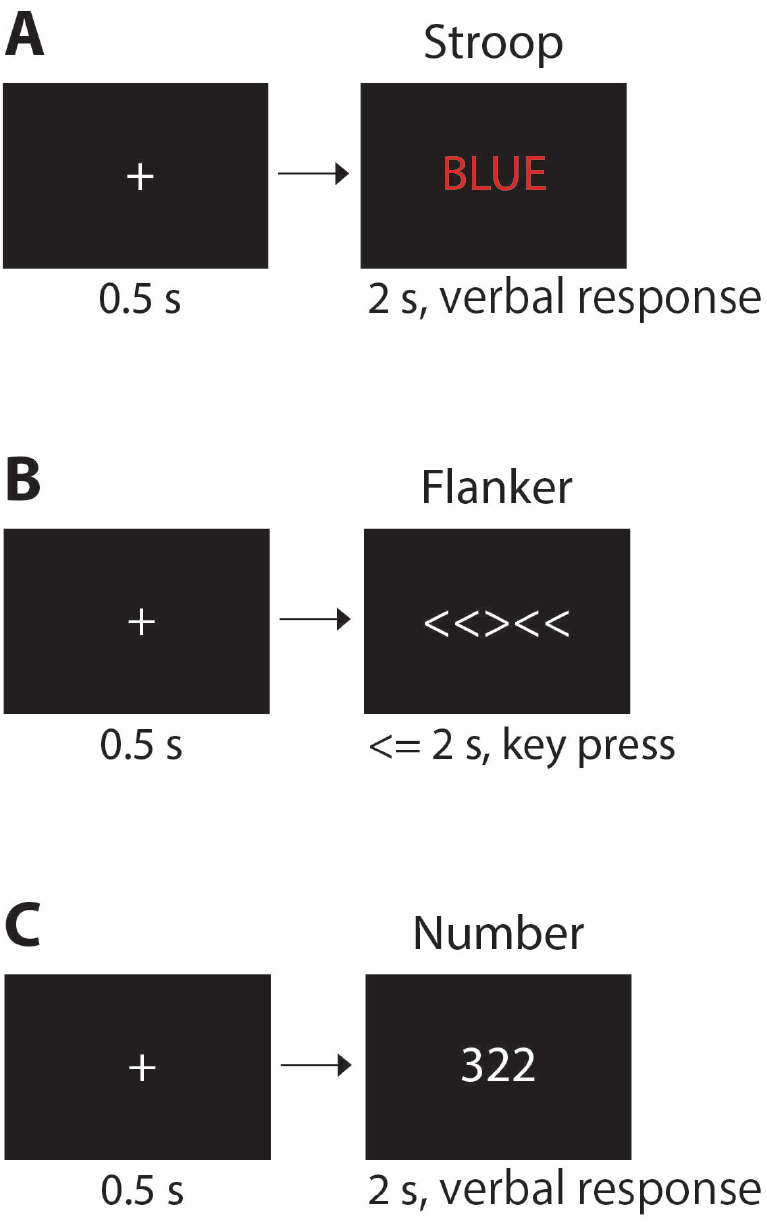
Experimental paradigms. Subjects performed the Stroop (A), Flanker (B), and Number (C) tasks in one session during intracranial neurophysiological recordings with depth electrodes. A standard session contained 18 blocks and each block comprised 30 trials of each task. (A) The Stroop task requires subjects to say the font color. In the congruent condition, the semantic meaning coincides with the font color, while the two conflict in the incongruent condition. (B) The Flanker task requires subjects to press the left or the right key to indicate the direction of the central arrow. In the congruent condition, all the arrows point in the same direction while in incongruent condition, the arrow in the middle points oppositely from the others (flankers). (C) The Number task requires subjects to say the position (“one”, “two”, or “three”) where the unique number is located. In the congruent condition, the target number and its position are the same while in the incongruent condition these are different. All trials in this figure show incongruent conditions.

The behavioral signature of this family of tasks is longer reaction time (RT) for incongruent stimuli (containing conflict) compared with congruent stimuli (conflict-free). For example, in the Stroop task, subjects take longer to name the font color of the word red when shown in green or blue font compared to red font. The increase in reaction time during incongruent conditions is due to interference from irrelevant but conflicting information and the selection among competing motor plans (Goghari and MacDonald, 2009; Miller and Cohen, 2001; Stroop, 1935).

Multiple studies have examined brain signals associated with each one of these cognitive control tasks, including measurements derived from human neuroimaging (Bunge et al., 2002; Bush and Shin, 2006; Coulthard et al., 2008; Fan et al., 2003; Robertson et al., 2014; Sani et al., 2021), human scalp electroencephalography (Hanslmayr et al., 2008; Janssens et al., 2018; Robertson et al., 2014), human invasive neurophysiology (Caruana et al., 2014; Koga et al., 2011; Oehrn et al., 2014; Sheth et al., 2012; Tang et al., 2016), and monkey neurophysiology (Blackman et al., 2016; Cole et al., 2009; Li et al., 2019; Nakamura et al., 2005). These studies have described an extensive network of frontal and parietal regions, and to a lesser extent temporal and other regions, that demonstrate distinct activation patterns between congruent and incongruent trials.

The neural differences between incongruent and congruent trials reported in those studies are often interpreted to reflect a correlate of a general and abstract notion of conflict irrespective of the specific combination of shapes, colors, input modalities, and behavioral outputs in any particular task. Thus, theories of cognitive flexibility refer to processes like error monitoring and conflict resolution which are thought to be independent of the specific sensory inputs that lead to such errors or conflict and which are also considered to be independent of the specific motor outputs involved in a particular task. For example, the conflict monitoring theory (Botvinick et al., 2001) expresses an abstract, domain-general notion of conflict: if certain neurons can detect the occurrence of conflict, these neurons should be activated regardless of the task implementation format. Such an abstract notion of conflict is also used in the interpretation of the Gratton effect, which describes the history-dependence of conflict modulation (Gratton et al., 1992; Sheth et al., 2012).

Combining these empirical and theoretical ideas, here we evaluate whether there are shared mechanisms involved in error monitoring and conflict resolution that are common across different sensory inputs and motor outputs. We focus on how conflict is represented in the brain by directly comparing neurophysiological responses during three cognitive control tasks, analyzing intracranial field potentials from 694 electrodes implanted in patients with pharmacologically intractable epilepsy. We hypothesize that conflict-related responses should show invariance to the stimulus properties within each task (within-task invariance). For example, in the Stroop task, we would expect that neural responses would distinguish congruent (RED/red, GREEN/green, or BLUE/blue) from incongruent (RED/green, RED/blue, GREEN/red, GREEN/blue, BLUE/red, or BLUE/green) conditions, irrespective of the specific font/semantic combination. Extending this hypothesis of within-task invariance to the comparisons across tasks, the assumption of an abstract notion of conflict led to our second hypothesis, that neural responses would distinguish conflict irrespective of whether incongruency is dictated by color, shape, or number stimuli, and also regardless of the specific motor outputs involved (across-task invariance). The results are consistent with the first hypothesis; neural signals that show modulation between incongruent and congruent trials are invariant to stimulus attributes within a task. In contrast, our results are inconsistent with the second hypothesis. The majority of the neural responses demonstrate robust modulation between incongruent and congruent trials that is task-specific and does not generalize across tasks.

## Results

We recorded intracranial field potentials (IFPs) from 16 epilepsy patients implanted with depth electrodes (**Table S1**). Subjects performed three cognitive control tasks: Stroop, Flanker, and Number (**Methods**, **Figure 1**). Importantly, subjects performed the three tasks during the same session, therefore enabling direct comparisons among the tasks. Each task began with a fixation cross shown for 500 ms at the center of the screen. The Stroop task stimulus consisted of color words (”RED,” “GREEN,” “BLUE,” or the corresponding traditional Chinese characters for patients in Taipei, **Methods**) shown in red, green, or blue font. Subjects were instructed to name the font color (**Figure 1A**). Conflict arises when the font color does not match the meaning of the word on the screen. The Flanker task stimulus consisted of five arrows in a horizontal row, and subjects were asked to press the left or the right key to indicate the direction of the *center* arrow (**Figure 1B**). Conflict arises when the center arrow points in the opposite direction to the other four arrows. The Number task required subjects to say the *position* of the unique number (”one,” “two,” or “three”) among three numbers shown in a horizontal row (**Figure 1C**). Conflict arises when the position of the unique number does not match the actual number (e.g., number “3” in position “1” in the stimulus “322”). For all the tasks, congruent and incongruent conditions, as well as the stimulus dimensions (word, color, arrow direction, number identity), were randomly interleaved and counterbalanced.

### Subjects showed behavioral evidence of conflict in the three tasks

Subjects showed high accuracy in all three tasks (**Figure S1**): Stroop (congruent) = 96.8 ± 0.9%; Stroop (incongruent) = 90.1 ± 2.2%; Flanker (congruent) = 96.3 ± 2.8%; Flanker (incongruent) = 90.2 ± 2.9%; Number (congruent) = 96.6 ± 1.7%; Number (incongruent) = 90.4 ± 2.3% (mean±SEM). On average, performance was significantly higher in the congruent condition compared to the incongruent condition in all three tasks; this difference reached statistical significance in the Stroop task (p=0.007, two-sided permutation test; 10,000 iterations), but not in the Flanker (p=0.33) or Number (p=0.39) tasks. These observations are consistent with previous work (Bush and Shin, 2006; Davelaar and Stevens, 2009; Eriksen and Eriksen, 1974; MacLeod, 1991; Sheth et al., 2012; Stroop, 1935; Tang et al., 2016), and are mostly ascribed to a ceiling effect (Carter and van Veen, 2007). Accuracy was high in the three tasks, with few error trials to have enough power to distinguish incongruent from congruent trials statistically. We focus exclusively on correct trials for the remainder of the study.

A hallmark of conflict in cognitive control tasks is the longer reaction time associated with incongruent trials (**Figure 2A-C**). As demonstrated in previous work (Davelaar and Stevens, 2009; Sheth et al., 2012; Tang et al., 2016), reaction times were longer during incongruent trials for all three tasks (Stroop: 1,122±8 ms vs. 953±7 ms, p<0.001; Flanker: 875±11 ms vs. 722±9 ms, p<0.001; Number: 1,110±8 ms vs. 972±8 ms, p<0.001; mean±SEM, two-sided permutation test, 10,000 iterations). The longer reaction times during incongruent trials were also statistically significant at the individual subject level in the majority of cases (Stroop: 16/16 subjects; Flanker: 14/16 subjects; Number: 15/16 subjects). Subject number 4 showed no significant difference in the Flanker and Number tasks, but this subject completed only half of a standard session. Absolute reaction times differ across tasks because of the distinct response modalities (verbal or keypress), because of the different number of response options (2 or 3), and because of the different processing modalities (language or vision). Therefore, to assess the difficulty of each task, we computed the ratio of reaction times in incongruent versus congruent trials. There was no significant difference in difficulty among the three tasks (**Figure 2D**, p=0.16, non-parametric one-way ANOVA). In sum, behavioral results were consistent with previous work and demonstrated almost ceiling accuracy and a longer reaction time associated with incongruent trials across the three tasks.

**Figure 2.**
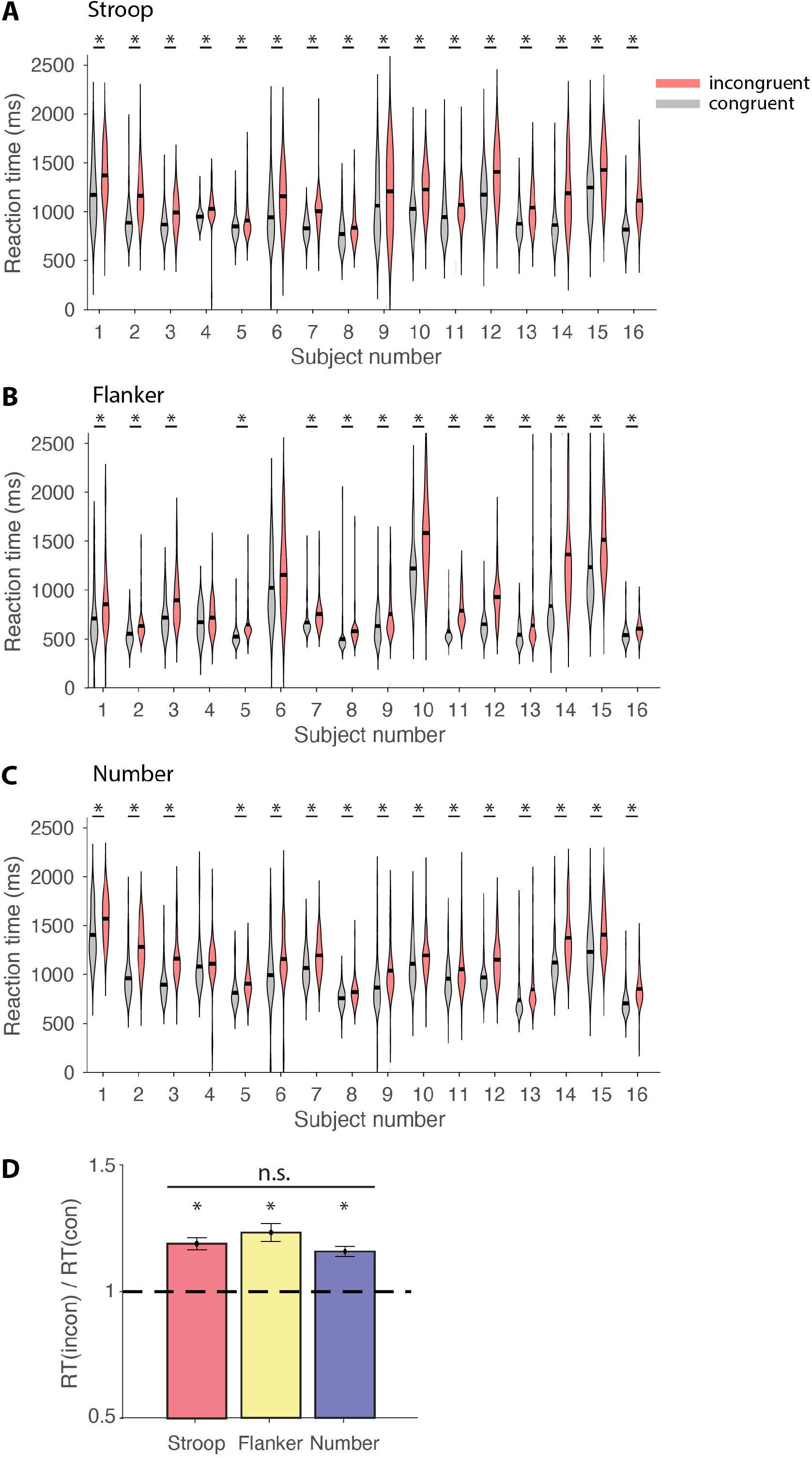
Subjects were slower in incongruent trials in the three tasks. **A-C**. Violin plots showing distribution of reaction times for each subject for congruent trials (gray) and incongruent trials (red) during the Stroop (**A**), Flanker (**B**), and Number (**C**) task. Only correct trials are shown. Black bars indicate mean reaction time. The asterisks denote statistically significant differences (two-sided permutation test, 10,000 iterations, α=0.05). **D**. There was no difference among the three tasks in task difficulty calculated as the ratio of reaction times during incongruent to congruent trials (one-way ANOVA, p=0.16). Asterisks denote significant differences in each task with respect to the null hypothesis corresponding to a ratio of 1 (two-sided permutation test, 10,000 iterations, α=0.05).

### Neural responses were modulated by conflict

We recorded intracranial field potential activity from 1,877 electrodes (**Table S1** reports the number of electrodes in each subject. We analyzed the activity from 694 bipolar-referenced electrodes that were not in the white matter (**Methods**); **Figure 3** and **Table S2** report the distribution of electrode locations. We focused on the neural activity in the theta band (4-8. Hz) because it constitutes a key component of cognitive control (Cavanagh and Frank, 2014; Gratton et al., 2018; Helfrich and Knight, 2016; Widge et al., 2019) and also on the high-gamma band (70-120 Hz) given its significance in sensory, motor, control, and other cognitive functions (Crone et al., 1998; Liu et al., 2009; Norman et al., 2019; Oehrn et al., 2014; Tang et al., 2016). Additional results in other frequency bands (alpha, beta, low-gamma) are reported in **Table S7**. In previous work (Tang et al., 2016), we reported that multiple electrodes showed activity in the high-gamma band that was modulated by the presence of conflict during the Stroop task. Consistent with previous work, **Figure 4** (left) depicts the high-gamma activity during the Stroop task of an electrode, located in the left orbitofrontal cortex, that showed enhanced responses during incongruent trials compared to congruent trials when aligning the neural signals to the behavioral response. The differences between incongruent and congruent trials were highly robust and could even be discerned in individual trials (compared **Figure 4C**, left versus **Figure 4B**, left). Notably, the enhancement associated with conflict was also evident when the neural responses were aligned to stimulus onset (**Figure 4D**, left).

**Figure 3.**
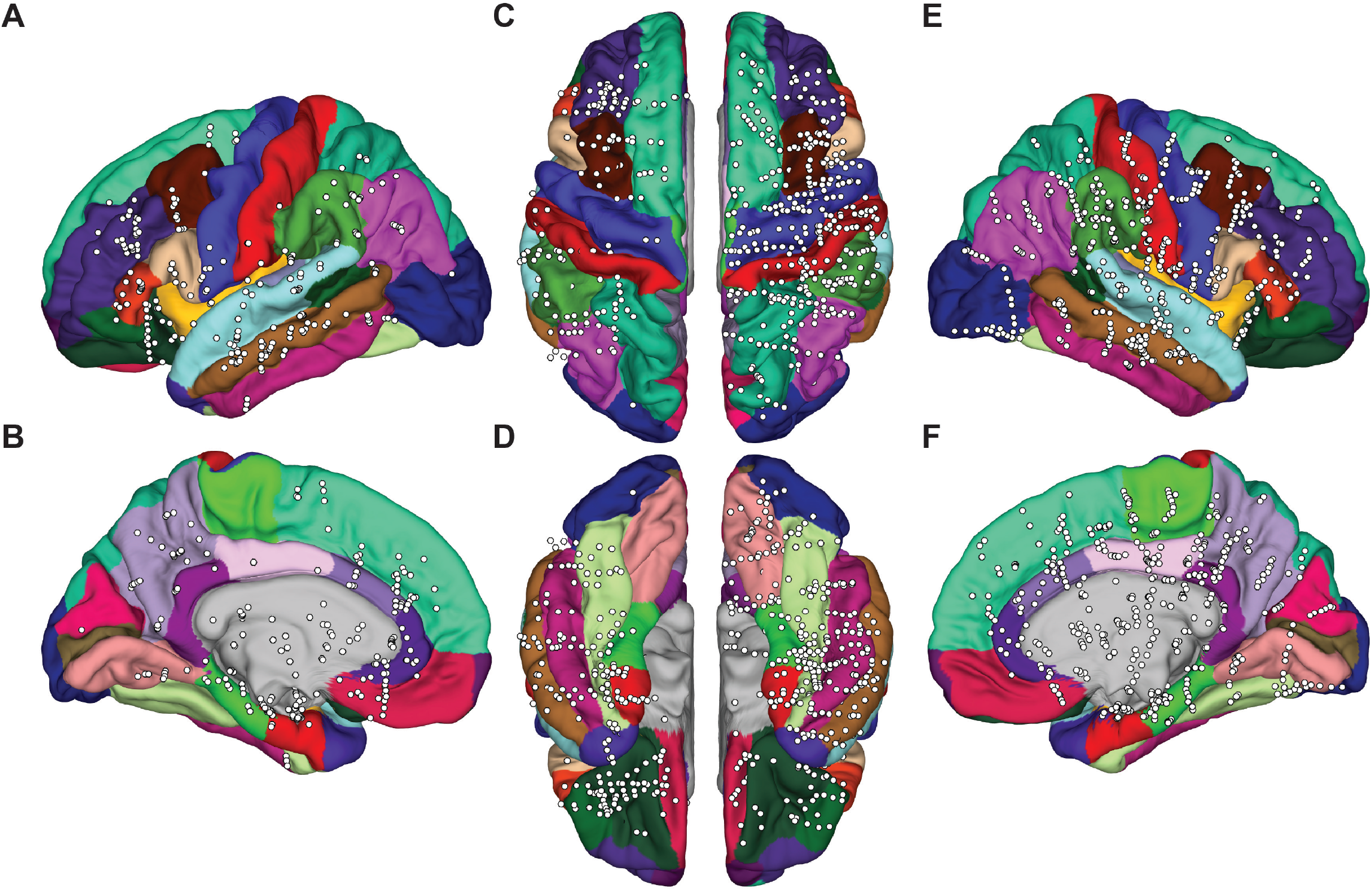
Electrode locations. Location of all n=694 electrodes overlayed on the Desikan-Killiany Atlas shown with different views. **A**: Left lateral; **B**: Left medial; **C**: Superior, whole brain; **D**: Inferior, whole brain; **E**: Right lateral; **F**: Right medial.

**Figure 4.**
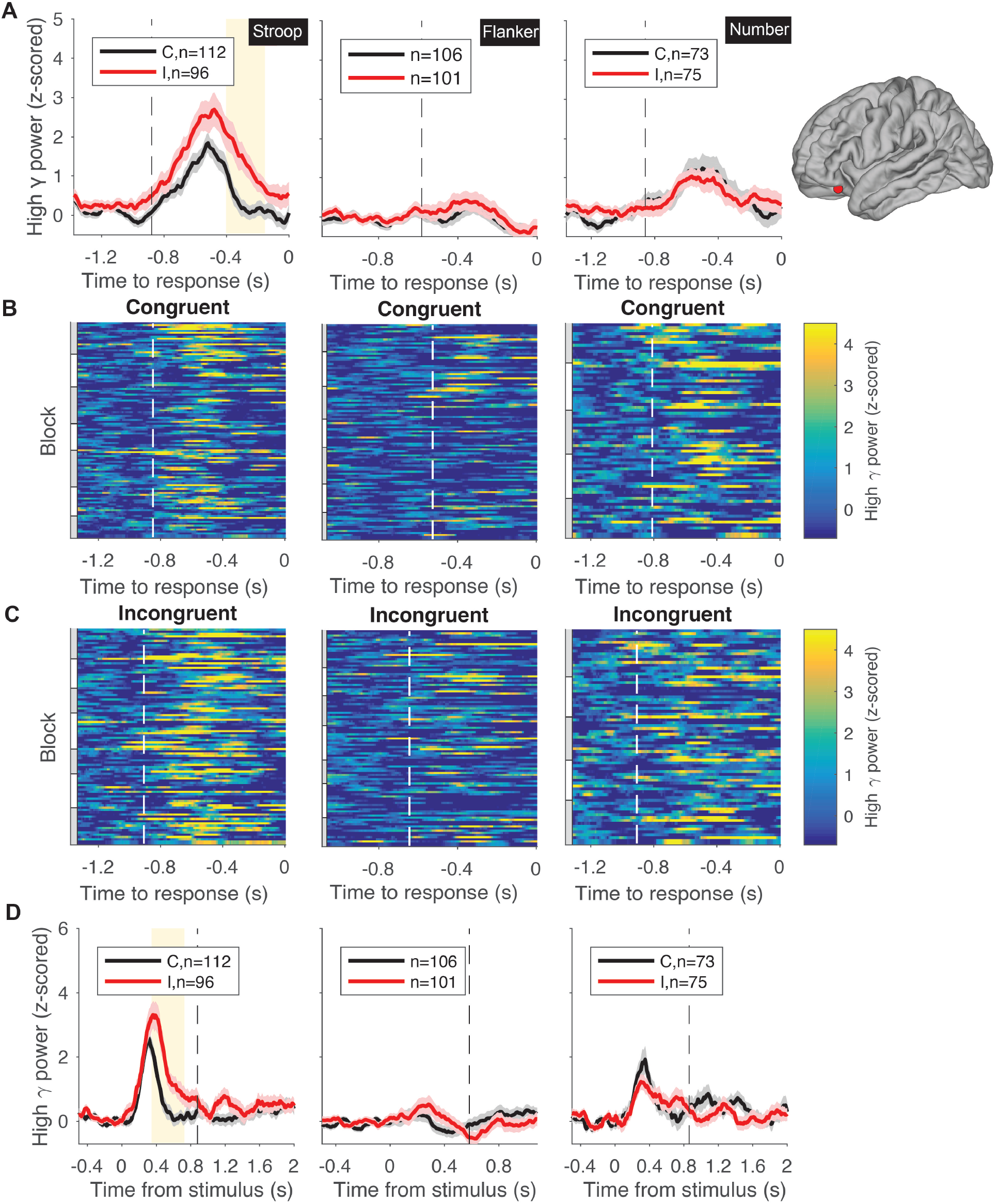
Example electrode in the left orbitofrontal cortex showing conflict modulation during the Stroop task. **A.** The traces show the mean±SEM z-scored high-gamma power aligned to behavioral response time for incongruent trials (red) and congruent trials (black) for each of the three tasks (Column 1: Stroop; Column 2: Flanker; Column 3: Number). The vertical dashed lines denote the average stimulus onset. Yellow background indicates statistically significant differences between congruent and incongruent trials (permutation test, 5000 iterations, α=0.05, **Methods**). The electrode location is shown on the right. **B-C**. Raster plots showing the neural signals in individual trials (see color scale on the right) for congruent (**B**) and incongruent (**C**) trials. The white dashed lines in the raster plots show the average stimulus onset time. These lines are shifted to the left in **C** compared to **B**, reflecting the longer reaction times during incongruent trials; Figure 2. Gray and white bars on the left of the raster plots represent different blocks. **C**. Z-scored high-gamma power (mean±SEM) aligned to stimulus onset. Vertical dashed lines denote the average behavioral response time. Yellow background indicates statistically significant difference between congruent and incongruent trials (permutation test, 5000 iterations, alpha=0.05, **Methods**).

An electrode was considered to be *conflict-modulated* if the band-filtered power during incongruent conditions was significantly different from that during congruent conditions for at least 150 consecutive milliseconds (permutation test, 5000 iterations, α = 0.05) *both* when responses were aligned to the behavioral response (**Figure 4A**, left) *and* also to the stimulus onset (**Figure 4D**, left, **Methods**). These strict selection criteria using both alignment to behavior *and* stimulus were implemented in order to exclude potential false positives. For example, signals from a visually responsive electrode could be confused for conflict modulation when aligning the neural responses to behavior due to the different reaction times between congruent and incongruent trials (**Figure 2A-C**). An example of such a visually responsive electrode located in the right lateral occipital cortex is shown in **Figure S2A-B**. Even though there seemed to be a difference between incongruent and congruent conditions when neural signals were aligned to the behavioral response (**Figure S2A**), this difference was absent when the neural signals were aligned to the stimulus onset (**Figure S2B**). Therefore, we do not consider this type of response to reveal any conflict modulation. Conversely, a motor responsive electrode could also be confused for conflict modulation when aligning the neural signals to stimulus onset for the same reasons (**Figure S2C-D**). Thus, the evaluation criteria for conflict modulation exclude purely sensory and purely motor responses.

**Figure 4** showed an example electrode that revealed conflict modulation in the high-gamma band during the Stroop task. Electrodes demonstrating robust conflict modulation were also observed during the Flanker and Number tasks. **Figure 5A** (middle) depicts the responses of an electrode in the right superior parietal that showed enhanced activity during incongruent trials in the Flanker task. As described for the Stroop task, conflict modulation was observed in single trials (**Figure S3A**, middle) and also when aligning the responses to stimulus onset (**Figure S3B**, middle). **Figure 5B** (right) depicts the responses of an electrode in the right precuneus that showed enhanced activity during incongruent trials in the Number task. **Figure S4A** (right) shows conflict modulation for this electrode during single trials and **Figure S4B** confirmed this conflict modulation even when aligning neural activity to the stimulus onset.

**Figure 5.**
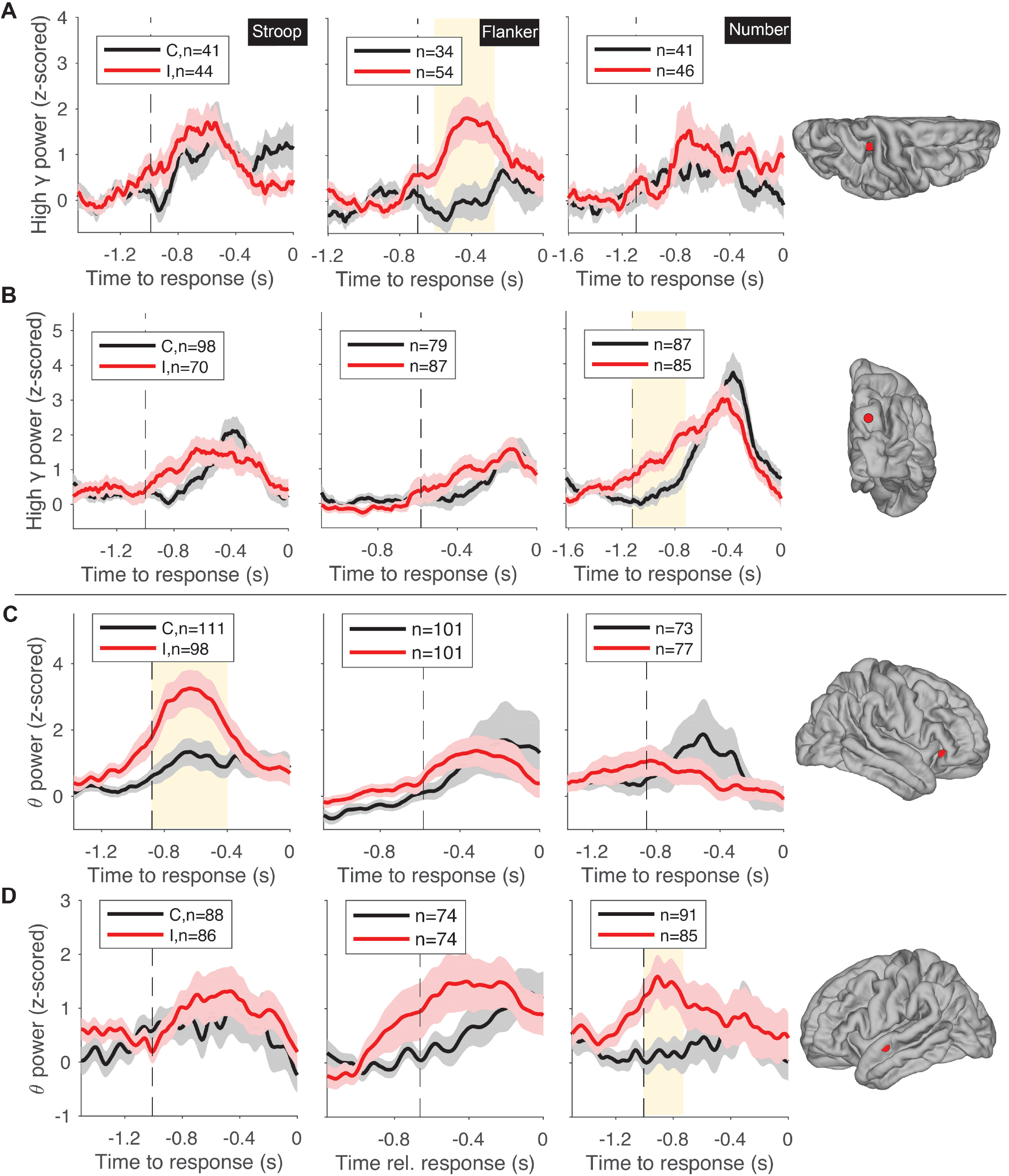
Example electrode showing conflict modulation in high-gamma (A-B) and theta band (C-D) during one task. (**A**: Flanker; **B**, **D**: Number; **C**: Stroop). The traces show mean±SEM z-scored high-gamma (**A-B**) or theta (**C-D**) power aligned to behavioral response time for incongruent trials (red) and congruent trials (black) for each of the three tasks (Column 1: Stroop; Column 2: Flanker; Column 3: Number). The vertical dashed lines denote the average stimulus onset. Yellow background indicates statistically significant differences between congruent and incongruent trials (permutation test, 5000 iterations, α=0.05, Methods). Electrode locations are shown on the right (**A**: right superior parietal; **B**: right precuneus; **C**: right pars triangularis; **D**: left superior temporal).

Similar results were observed when considering the theta frequency band. **Figure 6** shows an example electrode in the right precentral cortex that demonstrated conflict modulation in the theta frequency band during the Flanker task. Such modulation can be appreciated both in response-aligned signals (**Figure 6A**) and stimulus-aligned signals (**Figure 6D**) signals, as well as in individual trials (**Figure 6B-C**). **Figures 5C** and **5D** show example electrodes, in the right pars triangularis and left superior temporal cortex, that exhibited conflict modulation in the theta band for the Stroop and Number tasks, respectively. **Figures S5** and **S6** show responses in individual trials and stimulus-aligned signals for these two example electrodes.

**Figure 6.**
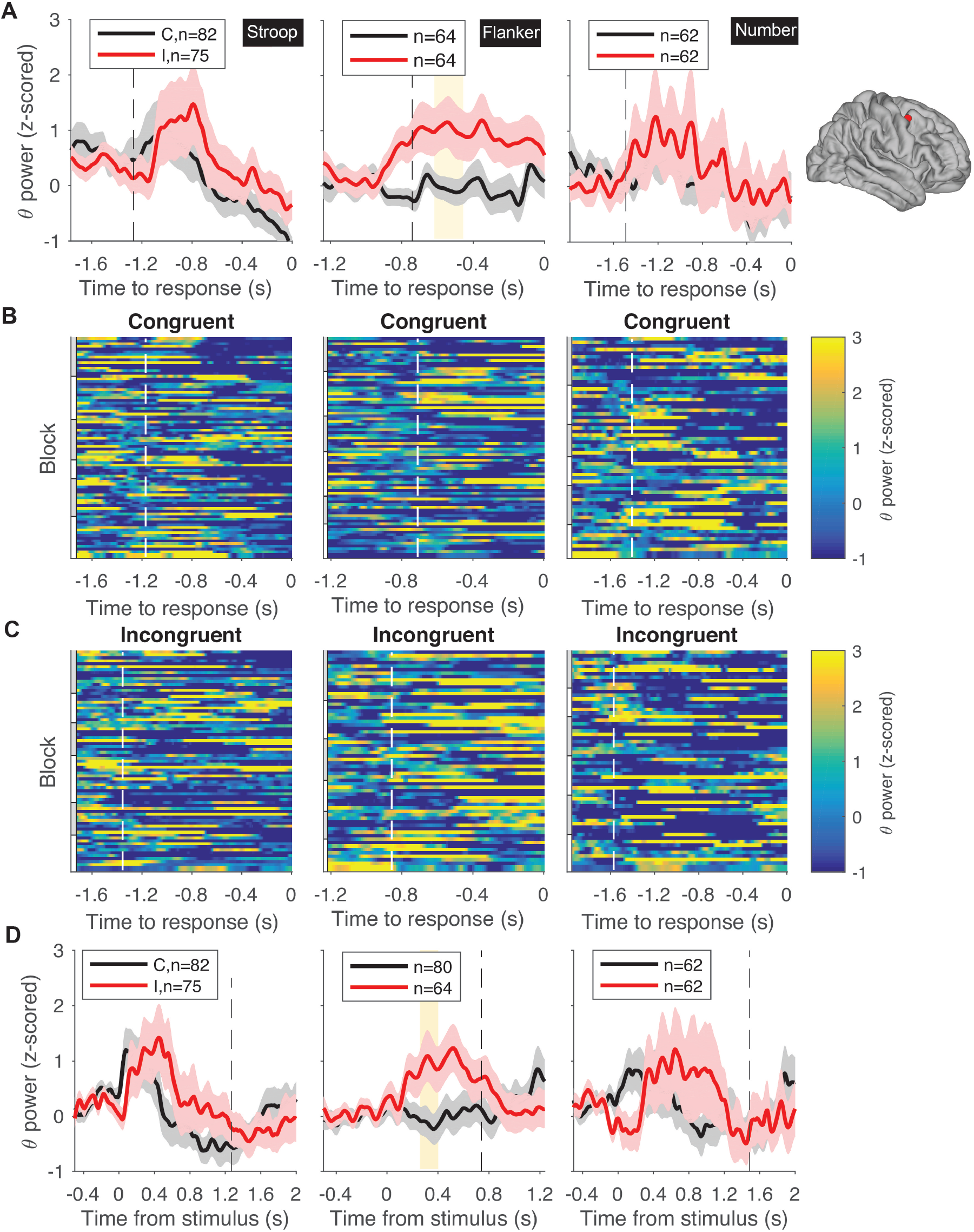
Example electrode in the right precentral cortex showing conflict modulation in the theta band during the Flanker task. **A**. The traces show the mean±SEM z-scored theta power (4-8 Hz) aligned to behavioral response time for incongruent trials (red) and congruent trials (black) for each of the three tasks (Column 1: Stroop; Column 2: Flanker; Column 3: Number). The format and conventions follow those in Figure 4. The vertical dashed lines denote the average stimulus onset. Yellow background indicates statistically significant differences between congruent and incongruent trials (permutation test, 5000 iterations, α=0.05, Methods). The electrode location is shown on the right. B-C. Raster plots showing the neural signals in individual trials (see color scale on the right) for congruent (**B**) and incongruent (**C**) trials. The white dashed lines in the raster plots show the average stimulus onset time. These lines are shifted to the left in **C** compared to **B**, reflecting the longer reaction times during incongruent trials (Figure 2). Gray and white bars on the left of the raster plots represent different blocks. **D**. Z-scored theta power (mean±SEM) aligned to stimulus onset. Vertical dashed lines denote the average behavioral response time. Yellow background indicates statistically significant difference between congruent and incongruent trials (permutation test, 5000 iterations, alpha=0.05, Methods).

Out of the total of 694 electrodes, we identified 134 electrodes (19%) that exhibited conflict modulation in at least one task in the high-gamma band (**Table S3**) and 109 electrodes (16%) when considering the theta band (**Table S4**). In most cases, conflict modulation was characterized by enhanced high-gamma-band power in the incongruent condition compared to the congruent condition, as illustrated in the three example electrodes shown in **Figure 4-6**. A few electrodes exhibited the reverse modulation direction where the congruent response was higher than the incongruent one (**Figure S11A**, middle). **Figure S7** shows the distribution of locations of electrodes revealing conflict modulation for each task. In sum, using strict criteria, we found electrodes that demonstrate robust conflict modulation in each of the three tasks, considering both high-gamma and theta band signals, evident in both behavior- and stimulus-aligned responses, and even in single trials.

### Neural signals in the high-gamma band during incongruent trials correlated with reaction times

Next we examined whether the neural signals were correlated with behavior. For each of the conflict-modulated electrodes, we plotted the mean high-gamma band power as a function of the reaction time (**Methods**). **Figure 7A** shows an example electrode located in the left rostral middle frontal cortex that was modulated by conflict during the Stroop task. The mean high-gamma power was not correlated with reaction times during congruent trials (**Figure 7A**, left, p=0.3), but there was a significant correlation during incongruent trials (**Figure 7A**, right, p=0.03). Similarly, **Figure 7B** shows an example electrode in the right superior frontal cortex that showed a correlation with reaction times during the Flanker task and **Figure 7C** shows an example electrode in the right inferior temporal cortex that showed a correlation with reaction times during the Number task. In all, 8.3%, 12.2%, and 10.2% of the conflict modulated electrodes showed a correlation with reaction time during incongruent trials, but not congruent trials, for the Stroop, Number, and Flanker tasks, respectively.

**Figure 7.**
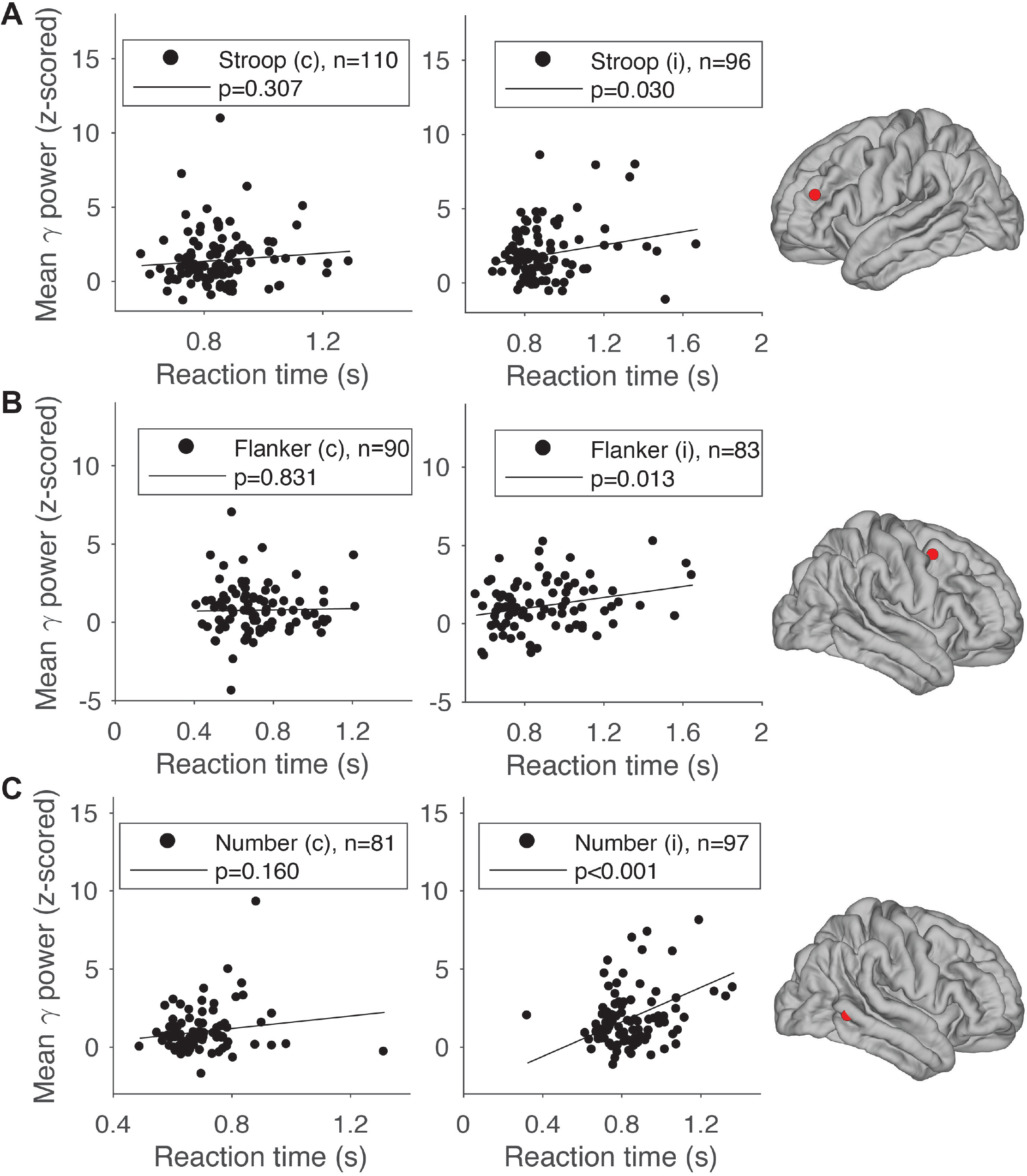
Example conflict modulated responses showing correlation with reaction time. The plots show the mean high-gamma power (z-scored) in each trial as a function of the reaction time for 3 example electrodes during the (**A**) Stroop (left rostral middle frontal cortex), (**B**) Flanker (right superior frontal cortex)), and (**C**) Number task (right inferior temporal cortex). Each point shows one trial. The number of trials is shown in each subplot. Electrode locations are shown on the right. The solid line shows the linear fits. Correlations were statistically significant for incongruent trials (right) but not congruent trials (left) (see p values in legend).

These observations did not extend to the theta band. Signals in the theta band showed a much weaker correlation with reaction times. We found only 4 conflict-modulated electrodes, 2 in the Flanker task, 2 in the Number task, and none in the Stroop task, that demonstrated a statistically significant correlation between theta band power and reaction times.

### Conflict representation exhibited within-task invariance

In each task, there are different stimuli that define conflict. For example, in the Stroop task, there are six different word/color combinations that are incongruent and three that are congruent (**Figure S8**). Our first hypothesis states that conflict modulation is *invariant* to the different stimuli defining incongruence within a task. To test this hypothesis, we evaluated whether the modulation of neural signals underlying conflict was evident only for certain stimuli defining incongruent trials but *not* other stimuli within each task, as opposed to a uniform modulation due to incongruent trials across the different stimuli within each task.

One might expect stimulus specificity given the extensive documentation of selective responses to different sensory inputs (e.g., (Liu et al., 2009), among many others). For example, an electrode located in visual cortical area V4 might be selective for color and respond differentially to RED compared to BLUE or GREEN. Indeed, consistently with previous work, we found multiple visually selective electrodes (Stroop: 15 electrodes; Flanker: 8 electrodes; Number: 0 electrodes; Total = 23 electrodes; **Methods, Table S5**). Similarly, we found 36 motor selective electrodes (Verbal response: 26 electrodes; Keypress response: 10 electrodes; Total = 36 electrodes; **Methods, Table S6**). Among these 23+36=59 electrodes, there were only 5 electrodes (3 visually-selective electrodes and 2 motor-selective electrodes and) that showed visual or motor selectivity *and* conflict modulation in the same task. These 5 electrodes constitute 8% of the visual/motor selective electrodes and 4% of all the electrodes that showed conflict modulation. Thus, the majority of electrodes that showed conflict modulation were *not* visually or motor selective.

To further investigate whether conflict modulation generalized across the different sensory inputs, we directly compared the responses to all possible stimuli within each task. **Figure S8A-I** describes the responses of an electrode in the left inferior parietal cortex for every word/color combination during the Stroop task. Conflict modulation cannot be ascribed to responses to specific word/color combinations; that is, conflict modulation showed *within-task invariance* with enhanced responses during incongruent trials for the six different possible incongruent word and font color combinations compared to the three different possible congruent word and font color combinations. An example in the theta band is shown in **Figure S8J-R**. Similarly, **Figure S9A-D** describes the responses of an electrode in the left orbitofrontal gyrus for every combination of central and peripheral arrow directions during the Flanker task. Conflict modulation in the Flanker task was also invariant within the task; that is, there was higher activity during both incongruent target/flanker combinations compared to the two congruent combinations. An example in the theta band is shown in **Figure S9E-H**. **Figure S10A-F** describes the responses of an electrode in the left superior frontal gyrus, showing that conflict modulation was evident for all the different incongruent conditions in the Number task. An example in the theta band is shown in **Figure S10G-L**.

To characterize the degree of within-task invariance at the electrode ensemble level, we used a machine learning decoding approach to assess whether we could decode the presence of conflict in individual trials (**Figure 8**). In all the decoding analyses, an SVM classifier with a linear kernel was trained after concatenating all the conflict modulated electrodes in each task. We used two neural features: the maximum and the mean band power during each trial (**Methods**), either for the high-gamma band (**Figure 8A**) or the theta band (**Figure 8B**). In all cases, we used cross-validation, separating the data into a training set and an independent test set and we randomly subsampled the data to ensure that the number of congruent trials matched the number of incongruent trials. To evaluate within-task invariance, the classifier was trained using only a subset of the different stimulus combinations and tested on different stimulus combinations. For example, in the first bars in **Figure 8A** and **8B**, the SVM classifier was trained with the neural responses to GREEN/red, GREEN/blue, BLUE/red, BLUE/green, GREEN/green, and BLUE/blue. The classifier’s performance was tested using the remaining conditions: RED/green, RED/blue, and RED/red. Even though the classifier was never exposed to the neural responses to any stimulus with the word “RED” during training, the classifier could extrapolate to identify conflict with those novel stimuli in the same task. Similar conclusions were reached for the other possible combinations of training and test stimuli within the Stroop task (**Figure 8**, red bars) and also for the different combinations in the Flanker (yellow bars) and Number (blue bars) tasks. In sum, both at the individual electrode level (**Figures S8-S10**) as well as at the electrode population level (**Figure 8**), and both in the high-gamma (**Figure 8A**) and theta band (**Figure 8B**), the results support the hypothesis that the neural signals modulated by conflict are largely independent of the specific combination of stimuli that give rise to incongruence *within* each task.

**Figure 8.**
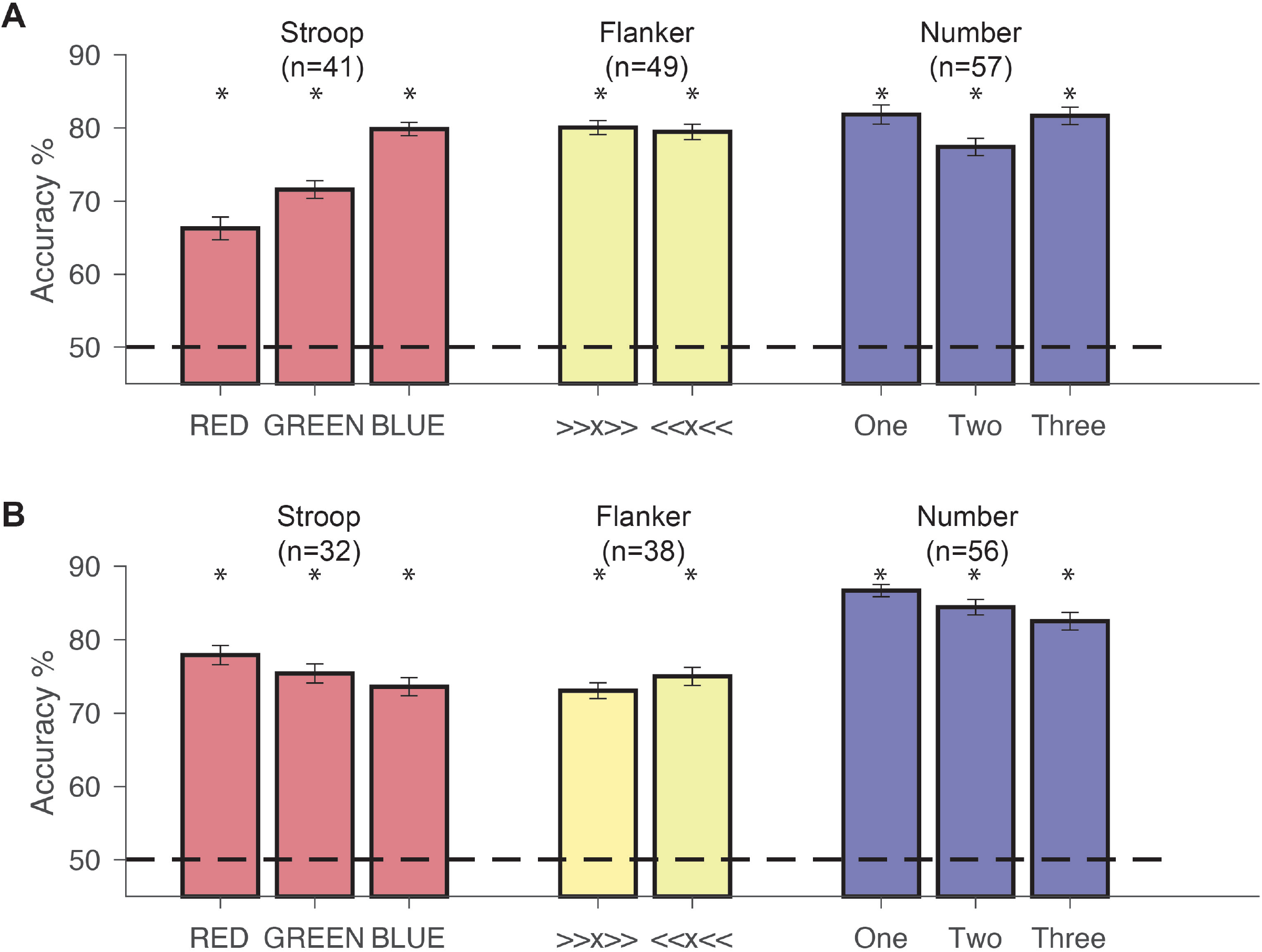
Within-task invariance in population-based decoding of conflict in single trials. Accuracy of support vector machine (SVM) classifier in discriminating incongruent from congruent trials extrapolating across conditions within each task (within-task invariance) using high-gamma (**A**) and theta (**B**) band power. **Stroop task.** The first bar labeled “RED” was trained using the “GREEN” trials (as in **DEF** in **Figure S8**) and “BLUE” trials (as in **GHI** in **Figure S8**), and tested on “RED” trials (as in **ABC** in **Figure S8**). A similar procedure was followed for the other combinations: in bar 2, the SVM was trained using “RED” and “BLUE” trials, and tested on “GREEN” trials. In bar 3, the SVM was trained using “RED” and “GREEN” trials, and tested on “BLUE” trials. **Flanker task.** In the first bar, the SVM was trained on “>>>>>” and “>><>>”, and tested on “<<<<<” and “<<><<”. In the second bar, training and testing data were reversed. **Number task.** In the first bar, the SVM was trained on trials where the correct answers were “two” (as in **CD** in **FigureS10**) or “three” (as in **EF** in **FigureS10**), and tested on trials where the correct answers were “one” (as in **AB** in **FigureS10**). Similarly in bar 2, the SVM was trained on “one” and “three”, and tested on “two”. In bar 3, the SVM was trained on “one” and “two”, and tested on trials whose target answers were “three”. For each task, the training and testing data for each condition were randomly subsampled to contain an equal number of congruent and incongruent trials. Error bars indicate s.e.m. over 50 sessions. The dashed line indicates chance performance (50%). Asterisks denote higher than chance accuracy (permutation test with 10,000 iterations, p<0.001 for all bars).

### Conflict-modulated electrodes were task-specific

Given the extrapolation across stimuli within a task, we next considered the hypothesis that neural signals representing conflict would also be independent of the specific sensory and motor characteristics of the task. We asked whether electrodes showing conflict modulation were *task-specific* (i.e., showing activity modulation during incongruent trials in *some* but not all tasks) or task-invariant (i.e., showing activity modulation during incongruent trials in *all* three tasks). The examples in **Figures 4-6** illustrate example electrodes with high specificity in their degree of conflict modulation. The electrodes in **Figure 4** and **Figure 5C** only revealed conflict modulation during the Stroop task (compare leftmost column to middle and rightmost columns). Similarly, the electrodes in **Figure 5A** and **Figure 6** showed conflict modulation only during the Flanker task (middle column), and the electrodes in **Figure 5B** and **Figure 5D** only during the Number task (rightmost column). These types of neural responses were representative of the majority of the data. Out of the total of 134 electrodes that showed conflict modulation in the high-gamma band, 118 electrodes (88%) exhibited modulation in one task but not in the other two tasks. Similarly, out of the total of 109 electrodes that showed conflict modulation in the theta band, 92 electrodes (84%) exhibited modulation in one task but not in the other two tasks. Although most electrodes demonstrated conflict modulation in one task only, there were 16 electrodes in the high-gamma band (12%) and 17 electrodes (16%) in the theta band that showed conflict modulation in two tasks. Three examples of electrodes that showed conflict modulation in two tasks are illustrated in **Figures S11-S12**. In **Figure S11A**, an electrode in the left inferior parietal cortex exhibited conflict modulation during the Stroop and Flanker tasks, but not during the Number task. Similarly, **Figure S11B** shows an electrode at the right supramarginal gyrus, demonstrating conflict modulation in the Stroop and Number tasks, but not during the Flanker task. **Figure S11C** shows an electrode in the insula exhibiting conflict modulation during the Flanker and Number tasks, but not during the Stroop task. These electrodes also showed conflict modulation when the neural signals were aligned to stimulus onset (**Figure S12**). **Table S3** and **Table S4** report the locations and task specificity for all the dual-task modulated electrodes for the high-gamma and theta band, respectively.

In sum, most electrodes demonstrated conflict modulation in one task and few electrodes showed conflict modulation in two tasks. Remarkably, we did not find *any* electrodes that were modulated by conflict in all three tasks. Given the complete absence of any task-invariant electrodes, we asked whether it is possible that we missed indications of invariance due to our stringent criteria. First, we considered whether it is possible that having elevated activity in the congruent condition could be a prerequisite to observe conflict modulation. The electrode in **Figure 4** showed conflict modulation for the Stroop task but not in the other two tasks. During the Flanker task, this electrode showed no elevated response whatsoever, and during the Number task, there was a high response with respect to baseline starting about 0.7 seconds before the behavioral response, but this increase was very much the same for congruent and incongruent trials. Thus, there can be activation in the congruent condition without conflict modulation. Similarly, in the example electrode in **Figure 5B**, there was an elevated response in the congruent condition during all three tasks. However, conflict modulation occurred only in the Number task. In total, 302 electrodes showed elevated high-gamma band responses during the congruent condition in at least one task. Among these electrodes, only 80 (26%) also showed conflict modulation. Moreover, the majority of these electrodes (70 out of 80) did not share the same task specificity, i.e., tasks showing conflict modulation did not match tasks displaying responses during the congruent conditions. In sum, multiple electrodes responded during the congruent condition without conflict modulation and multiple electrodes showed conflict modulation only in some task(s) while still showing a response during the congruent condition in other task(s). Thus, an elevated response during the congruent condition is neither necessary nor sufficient to show evidence of conflict modulation. Lack of task invariance cannot be attributed to lack of a response during the congruent condition.

Second, we asked whether lack of invariance could be attributed to the behavioral performance by subjects in a given task. In an extreme hypothetical, if a subject performs the Stroop task correctly and closes their eyes during the other two tasks, we might be misled into thinking that an electrode showed task-specificity. Several pieces of evidence argue against this possibility. First, most subjects showed high accuracy in *all* tasks, except subjects 1 and 6, who performed slightly less well on the Number and Flanker task, respectively (**Figure S1**). However, both subjects have electrodes that were modulated by either the Flanker or Number task, or both. Second, almost all subjects showed a clear behavioral conflict effect in the three tasks (**Figure 2A-C**), except for subject 4 who performed half of the sessions and did not show behavioral conflict in either the Flanker or Number task (**Figure 2B-C**). *All* subjects experienced conflict during the Stroop task at the behavioral level (**Figure 2A**). However, electrodes like the ones shown in **Figure 5ABD** and **Figure 6** clearly demonstrated no evidence of conflict modulation in the Stroop task. Finally, and even more conclusively, there were many examples of different electrodes in the same subject showing task-specificity for conflict modulation in *different tasks*. Therefore, lack of task invariance cannot be ascribed to cases where subjects showed conflict modulation in one task but not others at the behavioral level.

Third, the results presented thus far focus on the high-gamma and theta frequency bands. Although different frequency bands of intracranial field potential signals tend to be correlated (Bansal et al., 2012), it is conceivable that some of the electrodes may reveal task invariance in conflict modulation in other frequency bands. To evaluate this possibility, we repeated all the analyses in the following frequency bands **(Methods)**: alpha (8-12 Hz), beta (12-35 Hz), and low gamma (35-70 Hz). **Table S7** reports the number of electrodes that showed conflict modulation for each task and for each frequency band. Summarizing **Table S7**, we found conflict modulation in all frequency bands, though the total number of electrodes that showed modulation was highest in the high-gamma band. Consistently with the results described in the previous sections, the vast majority of electrodes revealed conflict modulation only in one task: alpha, 88%; beta, 94%; low-gamma, 93% (cf. 88% for the high-gamma band and 84% for the theta band). In all frequency bands, we observed a small fraction of electrodes that showed conflict modulation in two tasks. Importantly, we did not find any electrode that showed task-invariance in any of these other frequency bands.

Finally, the results presented thus far relied on highly rigorous pre-processing through bipolar referencing and stringent selection criteria by requiring a long window of 150 ms to identify significant differences between incongruent and congruent trials and 5,000 iterations of a permutation test. We relaxed all of these constraints by using global referencing, by evaluating a shorter duration threshold of 50 ms, and using a t-test. We analyzed 748 electrodes (this is more than the 694 electrodes reported so far because bipolar referencing reduces the number of electrodes). Using these more liberal criteria, we found two electrodes in the high-gamma band that showed task invariance, one located in the left superior frontal gyrus and the other one in the right rostral middle frontal gyrus. One of these electrodes is shown in **Figure S13**. Although the neural signals from this electrode are less compelling than the examples showing task modulation in a single task or two tasks (e.g., **Figures 4-6**), these observations hint at the possibility of a weaker and less localized signal common across tasks. Yet, even under these liberal selection criteria, only 0.3% of the total number of electrodes that we studied demonstrated task invariance in the high-gamma band and none in the other frequency bands.

In sum, the observations show that most of the electrodes reveal conflict modulation in only one task, and few electrodes show conflict modulation in two tasks. These results lead us to reject our second hypothesis of task invariance in cognitive control at the level of individual electrodes.

Electrode population level responses revealed task-specific conflict modulation in individual trials It is conceivable that individual electrodes could show task specificity while an ensemble of multiple electrodes might reflect task invariance. To evaluate this possibility, we investigated whether we could decode the presence of conflict at the *electrode population level* in individual trials, following the same procedure described in **Figure 8**. Depending on the specific question about task-specificity, each calculation used different combinations of training and test sets, as described below.

First, we asked whether the population of electrodes modulated by one task could classify the presence of conflict on another task. In **Figure 9**, we trained nine different classifiers using the high-gamma (**Figure 9A**) and theta (**Figure 9C**) band power. The first three bars used Stroop-only electrodes (like the ones in **Figure 4** and **Figure 5C**), the middle three bars used Flanker-only electrodes (like the ones in **Figure 5A** and **Figure 6**), and the last three bars used Number-only electrodes (like the ones in **Figure 5B** and **Figure 5D**). The classifiers were trained and tested using cross-validation using only responses from the Stroop task (red), using only responses from the Flanker task (yellow), or using only responses from the Number task (blue). The Stroop-only electrode population yielded significant classification performance when trained and tested on the Stroop task (permutation test, 10,000 iterations, one-sided, p<0.001, **Figure 9A** and **9C**, bar 1), and the Flanker-only population yielded significant classification performance when trained and tested on the Flanker task (p<0.001, **Figure 9A** and **9C**, bar 5). The population of Number-only electrodes yielded significant classification performance when trained and tested on the Number task (p<0.001, **Figure 9A** and **9C**, bar 9), but also when trained and tested on the Stroop task (p<0.01, **Figure 9A** and **9C**, bar 7). Although the Number-only electrode population could detect conflict in the Stroop task, the performance on the Number task was still significantly higher than that on the Stroop task (p<0.001).

**Figure 9.**
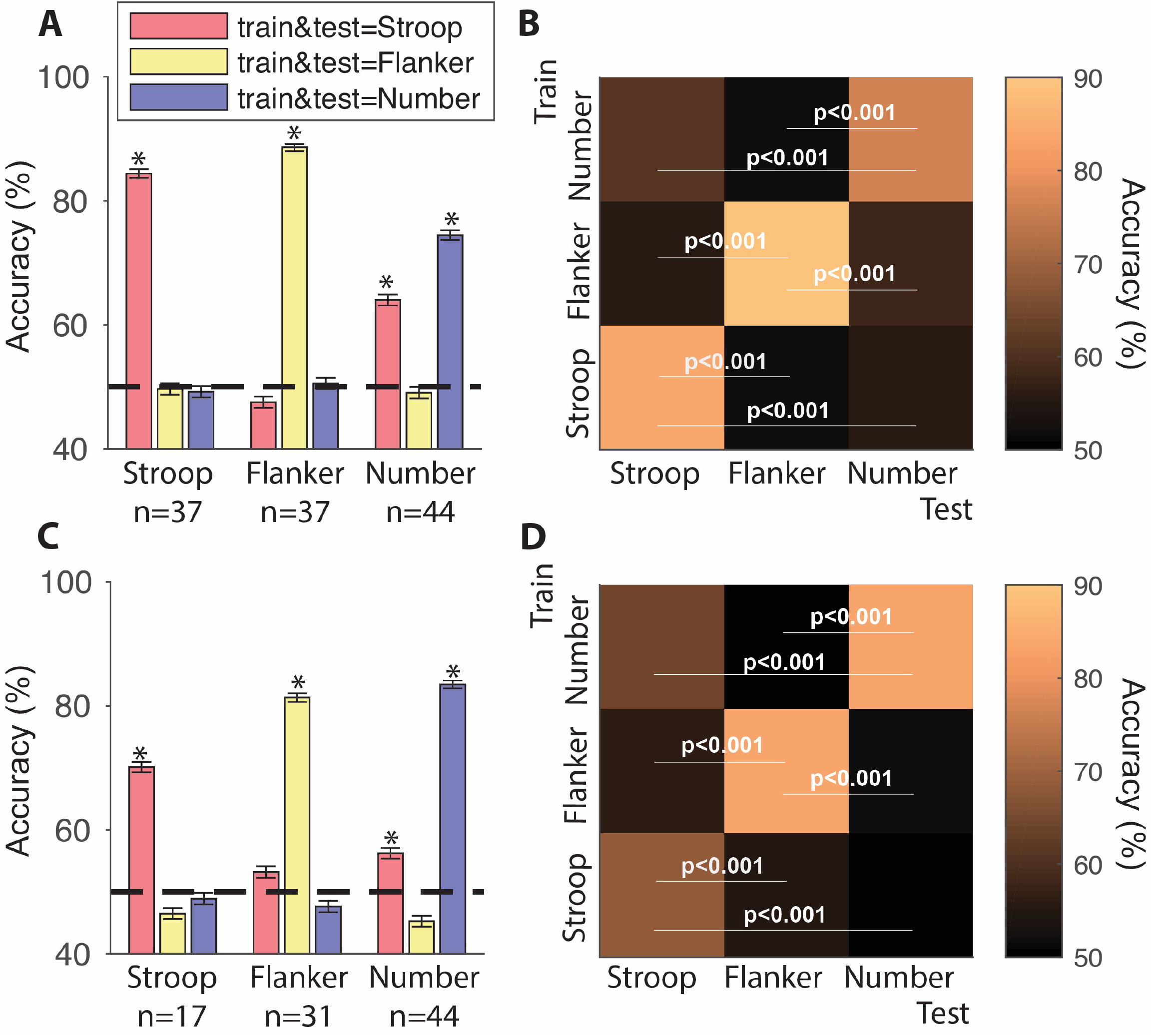
Task-specificity in population-based decoding of conflict in single trials. **A, C**. Accuracy of SVM classifier in congruent/incongruent discrimination when using a population of Stroop-specific electrodes (first three bars, n=37), Flanker-specific electrodes (next three bars, n=37), or Number-specific electrodes (last three bars, n=44). The SVM classifier was trained and tested with ten-fold cross-validation x 50 sessions of random sampling of trials using high-gamma (**A**) and theta (**C**) band power from the Stroop (red), Flanker (yellow), or Number (blue) tasks. Asterisks indicate that performances are significantly higher than chance (permutation test, 10,000 iterations, one-sided, p<0.001). **B, D**. Cross-task training and testing. Here we used the same three populations from part **A** and **C.** The SVM classifier was trained on one task and tested on the other two tasks. The diagonal corresponds to training and testing within the same task and the off-diagonal entries show cross-task extrapolation. P values indicate the comparison between within-task and cross-task testing performances in each electrode population (permutation test with 10,000 iterations, one-sided). Accuracy is reflected by the color of each square (see color map on right).

We next sought to assess whether a classifier trained only with data from one task *could extrapolate to detect conflict in a different task*. We trained three classifiers, one using data from the Stroop task only, one using data from the Flanker task only, and one using data from the Number task only using high-gamma (**Figure 9B**) and theta (**Figure 9D**) band power. Then we tested each classifier with data from the Stroop, Flanker, and Number tasks, respectively. We performed this analysis using Stroop-only, Flanker-only, and Number-only electrodes. Note that this is different from the analyses in **Figure 9A** and **9C**, where the training and test data were always from the *same task* for each classifier, whereas here, the training and test data can be from different tasks. Here, even when the classifier is trained and tested using data from the same task, we still performed cross-validation across trials to avoid overfitting. The results of these cross-task classification analyses are presented in **Figure 9B**. As expected, the highest classification accuracies were observed for *within-task training and testing* (diagonal tiles in **Figure 9B** and **9D**). These three conditions not only exhibited better than chance accuracies (high-gamma: 82.6 ± 6.9%, theta: 79.5 ± 8.2%), but also significantly higher performance than all the corresponding cross-task accuracies (high-task: 55.3 ±4.3% **Figure 9B**, p<0.001; theta: 53.6 ±5.0% **Figure 9D**, p<0.001). In sum, even at the electrode population level, we observed minimal ability to detect conflict when a decoder was trained and tested in different tasks despite the fact that there was high accuracy in distinguishing conflict in individual trials within each task.

## Discussion

We studied the neural mechanisms underlying conflict resolution during cognitive control by recording intracranial field potentials from 694 electrodes in 16 subjects who performed three different tasks: Stroop, Flanker, and Number (**Figure 1**). Subjects showed increased reaction times during incongruent trials compared to congruent trials (**Figure 2**), a hallmark of cognitive control (Bush and Shin, 2006; Eriksen and Eriksen, 1974; Mayr et al., 2003; Stroop, 1935). Consistent with previous studies (Caruana et al., 2014; Gaetz et al., 2013; Koga et al., 2011; Tang et al., 2016), we found robust modulation of neural signals in the high-gamma frequency band when comparing incongruent versus congruent trials (**Figures 4, 5**). Conflict modulation was also present in the theta band (**Figures 5, 6**) and other frequency bands (**Table S7**). Modulation was evident both when aligning neural signals to the behavioral responses and to stimulus onset (**Figures 4, 6, S3-6, S12**), could be appreciated even in single trials (**Figures 4**, **6**, **Figures S3-S6**), and showed *within-task invariance* to the different combinations of visual inputs (**Figures 8, S8-S10**). Surprisingly, despite this robust within-task invariance, most of the electrodes showed *task-specificity*, with clear incongruent/congruent modulation in only one task but not in the other two (**Figure 4-6**). A few electrodes showed task-modulation in two tasks but not the third task (**Figure S11**).

We were concerned about multiple potential factors that could masquerade as task specificity. First, if subjects failed to perform a given task adequately or failed to show conflict at the behavioral level in a given task, one may wrongly conclude that there is task specificity in the neural responses. However, all subjects revealed high accuracy in the three tasks (**Figure S1**), and the vast majority of subjects showed conflict at the behavioral level (**Figure 2A-C**). We observed task-specificity in the neural signals even in subjects that showed conflict at the behavioral level in all three tasks. Furthermore, there were electrodes in the same subject with different task-specificity, ruling out an explanation of task-specificity based on poor behavior. Another potential behavioral factor that could correlate with task-specificity would be if one task showed more “conflict” than the other tasks. However, the reaction times showed that the three tasks were comparable in terms of their degree of behavioral conflict (**Figure 2D**). Additionally, if there were a gradient of “conflict,” say Stroop > Flanker > Number, one may expect to observe task-specificity only for Stroop, followed by electrodes that show modulation in both Stroop and Flanker, followed by electrodes that show invariance to all three tasks. However, this possibility does not match our observations since we report task-specific electrodes for each of the three tasks.

We used stringent criteria to ascribe conflict modulation to an electrode (**Methods**). We considered whether it is possible that task-specificity could be dictated by a lack of response during congruent trials, but this was not the case. We showed that an elevated response during the congruent condition was neither necessary nor sufficient to show evidence of conflict modulation. We also considered whether our strict selection criteria could have biased the interpretation of our results. However, an inspection of the individual trials reveals robust modulation during incongruent trials compared to congruent trials in some tasks but not others (**Figures 4, 6, S3-6**). Even after relaxing pre-processing by using a global reference and a much less stringent statistical threshold, only two electrodes (less than 1%) revealed task invariance. Similar conclusions were reached when examining other frequency bands (**Table S7**).

We also considered an electrode ensemble machine learning decoding approach (**Figures 8-9**). Population-based decoding is highly sensitive and could, in principle, uncover a task-invariant representation even if we mainly observe specificity in individual trials and individual electrodes. However, the decoding results also support the conclusion of clear within-task invariance (**Figure 8**) and a largely task-specific representation (**Figure 9**). These decoding results cannot be ascribed to drifting neural signals or non-stationarities in the data. First, previous work showed that intracranial field potentials tend to be very stable within a session, and even across recording sessions spanning multiple days (Bansal et al, 2012). Second and most importantly, the total of 18 blocks with different tasks were randomly interleaved and the signals were still more consistent within a task than across tasks.

It is important to emphasize that our sampling of brain locations is extensive but certainly not exhaustive (**Figure 3**). Therefore, it is quite possible that there are other brain regions that represent conflict in a task-invariant fashion in areas that we could not sample here. It is also relevant to highlight that our study focuses on intracranial field potentials; these signals combine the activity of large numbers of neurons. It is conceivable that individual neurons might show more, or less, task invariance than the results reported here. However, recent studies examining single unit activity in the frontal cortex are also consistent with a lack of task-invariance in cognitive control (Ebitz et al., 2020; Fu et al., 2019; Smith et al., 2019). Several studies have shown that frontal cortex neurons demonstrate “mixed selectivity” (Rigotti et al., 2013). Such mixed selectivity is a good summary of the results described here at the level of intracranial field potentials, which seem to reflect a combination of conflict and task-specific demands.

These task-specific electrodes were located in multiple regions within the frontal, parietal, and temporal lobes, and to a lesser degree, the occipital lobe. Several studies documented differential activation during incongruent versus congruent trials in the frontal cortex (Bunge et al., 2002; Caruana et al., 2014; Fan et al., 2003; Milham and Banich, 2005; Parris et al., 2019; Robertson et al., 2014; Sheth et al., 2012; Tang et al., 2016). The locations shown in **Figure S7** and **Table S3-4** are also consistent with many studies documenting responses during cognitive control in the parietal cortex (Bunge et al., 2002; Bush and Shin, 2006; Coulthard et al., 2008; Fan et al., 2003), temporal cortex (Bush and Shin, 2006; Fan et al., 2003), occipital cortex (Egner and Hirsch, 2005; Fan et al., 2003; Janssens et al., 2018), and other brain areas such as the insula (Menon and Uddin, 2010). These results suggest that cognitive control processes recruit distributed and task-specific networks rather than a single brain region (Dosenbach et al., 2007; Dosenbach et al., 2006; Fan et al., 2003; Marek and Dosenbach, 2018).

Several previous studies reported signals that differ between incongruent and congruent trials at the level of individual neurons (Sheth et al., 2012; Smith et al., 2019), intracranial field potentials (Caruana et al., 2014; Koga et al., 2011; Oehrn et al., 2014; Tang et al., 2016), in scalp electroencephalography signals (Hanslmayr et al., 2008; Janssens et al., 2018; Robertson et al., 2014), and in functional neuroimaging signals (Bunge et al., 2002; Goghari and MacDonald, 2009; Milham and Banich, 2005; Parris et al., 2019). These signals have been interpreted and modeled as reflecting conflict (Heilbronner and Hayden, 2016; Ridderinkhof et al., 2004; Shenhav et al., 2013). Multiple potential confounds could lead investigators to infer that such signals represent task-invariant conflict, even though the signals only reflect task-specific differences between congruent and incongruent trials. The most obvious and ubiquitous reason for this inference is extrapolation based on studies that draw conclusions from a single task. With a single task, it is impossible to assess task-invariance in conflict modulation. It is possible to draw inferences about *potential* invariance by comparing results in different studies; however, precise anatomical comparisons across subjects can be challenging, especially when considering coarse signals that smooth over large numbers of neurons. Inferences across studies do not necessarily imply that the same neural circuits represent conflict in an abstract format. Another potential confound is the distinction between signals aligned to the stimulus or to the behavioral response, which requires a careful comparison of the temporal dynamics of the neural responses. Stimulus-specific neural signals could be misconstrued as conflict modulation if neural responses are aligned to the motor output (e.g., **Figure S2AB**), and motor-specific neural signals could be misconstrued as conflict modulation if neural responses are aligned to the stimulus onset (e.g., **Figure S2CD**). Thus, either due to single-task studies or spatial and temporal averaging, many previous studies likely reflect task-specific modulation rather than an abstract conflict signal.

We deliberately designed the tasks to be different in terms of the sensory inputs and motor outputs. Conflict relies on a discrepancy between color and semantic meaning (Stroop), comparison between shapes (Flanker), and the meaning of numbers and positional encoding (Number). Subjects used either verbal responses (Stroop, Number) or keypress responses (Flanker) as output. We conjectured that a general, abstract, signature of cognitive control should be independent of the inputs and outputs that define conflict. However, we found no such task-invariant conflict signals. It is tempting to assume that neural signals from electrodes that show conflict modulation in two tasks (e.g., **Figure S11**) correlate with the common aspects of two tasks. For example, electrodes that showed modulation exclusively during the Stroop and Number tasks (e.g., **Figure S11B**) might be involved in conflict expressed through verbal output. However, caution should be exercised in this type of interpretation because we did not test the Flanker task using a verbal output. The majority of electrodes responded in a task-specific manner, arguably demonstrating engagement in conflict only through specific sensory-motor combinations. Collectively, our results indicate that cognitive control is orchestrated by largely distinct and distributed networks dictated by the specific demands of each task.

## Materials and Methods

### Subjects and recording procedures

Subjects were 16 patients (8 female, ages 12-62, **Table S1**) with pharmacologically intractable epilepsy treated at Taipei Veterans General Hospital (TVGH), Boston Children’s Hospital (BCH), Brigham and Women’s Hospital (BWH), and Johns Hopkins Medical Hospital (JHMH). The electrode locations are purely dictated by clinical considerations, precluding any quantitative estimation of sample size at study design. The target number of subjects, 16, was decided during study design based on historical data of electrode distributions from previous studies. This study was approved by the institutional review board in each hospital and was carried out with subjects’ informed consent. Subjects were implanted with intracranial depth electrodes (Ad-Tech, Racine, WI, USA). The electrode locations were dictated by the clinical needs to localize the seizure focus in each patient (Fried et al., 2014). The total number of electrodes was 1,877 (**Table S1**). Neurophysiological data were recorded using XLTEK (Oakville, ON, Canada), Bio-Logic (Knoxville, TN, USA), Nihon Kohden (Tokyo, Japan), and Natus (Pleasanton, CA). The sampling rate was 2048 Hz at BCH and TVGH, 1000 Hz at JHMH, and 512 Hz at BWH. All data were bipolarly referenced, unless stated otherwise. There were no seizure events in any of the sessions. Electrodes in the epileptogenic foci, as well as pathological areas, were removed from analyses.

### Task procedures

Each subject completed three tasks in a single recording session: Stroop, Flanker, and Number. A schematic rendering of the tasks is shown in **Figure 1**. Each session contained 18 blocks, with 30 trials of one task (Stroop, Flanker, or Number) per block. The target number of trials was pre-defined based on the results of one of our previous studies (Tang et al, 2016) and based on a pilot study with 4 healthy volunteers where we confirmed that conflict (i.e., reaction time difference between congruent and incongruent trials) could be robustly detected with this number of trials. Per our IRB protocols, subjects can stop testing at any time; subjects who completed different numbers of blocks are indicated in **Table S1** in bold font. Subjects completed the tasks in normally lighted and quiet rooms. The experiments were written and presented using the Psychtoolbox extension in Matlab_R2016b (Mathworks, Natick, MA). Subjects viewed and completed the experiment using a 13-inch Apple Mac laptop. Stimuli subtended approximately 5 degrees of visual angle and were centered on the screen. Before each experiment started, each subject went over a short practice session until the instructions were fully understood. During the actual experiment, no correct/incorrect feedback was provided.

All trials started with 500 ms of fixation, followed by stimulus presentation. The stimulus was presented for 2,000 ms (Stroop, Number), or until the minimum of 2,000 ms and the subject’s key response time (Flanker). The stimuli were presented in white (Flanker, Number) or red/green/blue font color (Stroop), on a black background. For those subjects in Taipei, Stroop task stimuli were presented in traditional Chinese characters. Subjects provided a verbal response recorded using a Yeti microphone with an 8,192 Hz sampling rate (Stroop, Number), or a two-alternative keypress response using the left and right keys on the experiment laptop (Flanker).

### Electrode localization

We used the iELVis (Groppe et al., 2017) pipeline to localize the depth electrodes. Pre-implant MRI (T1, no contrast) was processed and automatically segmented by Freesurfer (Dale et al., 1999; Reuter et al., 2012), followed by co-registering the post-implant CT to the processed MR images. Electrodes were then identified visually and marked in each subject’s co-registered space using the BioImage Suite (Joshi et al., 2011). Each electrode was assigned an anatomical location (parcellated cortices (Desikan et al., 2006), white matter, subcortical regions, or unknown locations) using the Freesurfer localization tool. Unknown locations could be due to brain lesions or pathological brain areas. Electrodes in the white matter, ventricles, cerebellum, and unknown locations were excluded from analyses. Out of a total of 1,877 electrodes, we included 694 bipolarly referenced electrodes in the analyses. To show the position of electrodes from different subjects (**Figure 3 and S7**), electrode locations were mapped onto the MNI305 average human brain via an affine transformation (Wu et al., 2018).

### Behavioral analyses

The content of verbal responses (Stroop and Number tasks) was transcribed offline. The transcription was blind to the ground truth answers as well as neural responses. The behavioral reaction time for verbal response (Stroop and Number tasks) was determined as the first time the energy of the soundtrack was three standard deviations above the mean energy of the whole trial. Any noise (e.g., door slam, coughing, etc.) before the actual trial response was carefully identified and smoothed to prevent false automatic identification of behavioral response time. The keypress reaction time (Flanker task) was recorded by the Psychtoolbox code. Reaction times are shown in **Figure 2**.

### Preprocessing of intracranial field potential data

A zero-phase digital notch filter (Matlab function “filtfilt”) was applied to the broadband signals to remove the AC line frequency at 60 Hz and harmonics. For each electrode and each task, trials with amplitudes (max-min voltage from fixation onset to stimulus off) larger than three standard deviations above the mean amplitude across all trials were considered as containing artifacts and excluded from analysis (Bansal et al., 2012). The percentage of trials excluded by this criterion was 1.05% (Stroop), 1.25% (Flanker), and 1.29% (Number).

### Single electrode analysis of modulation by conflict

We computed the high-gamma band (70-120 Hz), low-gamma band (35-70 Hz), beta band (12-35 Hz), alpha band (8-12 Hz), and theta band (4-8 Hz) power of the intracranial field potential signals by using a multi-taper moving-window spectral estimation method implemented in the Chronux toolbox (Mitra and Bokil, 2008). The time-bandwidth product, number of tapers, and size of moving window used for each frequency band are listed in **Table S7** (Tang et al., 2016). Throughout the paper, we focus on the high-gamma band and theta band signals. The power in the corresponding frequency band was z-scored by subtracting the mean high-gamma power during the baseline period (500 ms before stimulus onset) and dividing by the standard deviation of the high-gamma power during the baseline. Only correct trials were included in the analyses.

First, we examined whether an electrode exhibited any response at all to the stimuli. An electrode was defined as “responsive” if the z-scored high-gamma power during the congruent condition was larger than 1 for at least 150 consecutive milliseconds (15 x 200 ms window shifted by 10 ms), starting from stimulus onset to average behavioral response time. To determine whether an electrode showed conflict modulation (**Tables S3**, **S4**, and **S7**), we compared the band power between the congruent and incongruent conditions of each task. For each time bin (200 ms shifted by 10 ms), we compared the band power of incongruent versus congruent trials using a permutation test with 5,000 iterations (*α* = 0.05). An electrode was denoted as showing conflict modulation if the following two criteria were satisfied: (1) The band power of incongruent trials was significantly different from the high-gamma power in congruent trials for at least 150 consecutive milliseconds (15 x 200 ms window shifted by 10 ms); (2) Criteria (1) was satisfied in both behavioral response-aligned and stimulus-aligned conditions (Tang et al., 2016). When the band power was aligned to behavioral response, selection criteria were applied to the time window starting from the *average stimulus onset* to the behavioral reaction time. When the band power was aligned to the stimulus, the time window was from stimulus onset to *average behavioral reaction time*.

An electrode was considered to be visually responsive if the maximum z-scored high-gamma band power was larger than 2 during the 300 milliseconds after stimulus onset. An electrode was considered showing motor responsive if the maximum z-scored high-gamma band power was larger than 2 during 300 milliseconds before behavioral response and the power continually increased during this time window. An electrode was considered to be visually selective for a particular task if it was visually responsive to stimuli only in one task. An electrode was deemed to show motor selectivity if it was motor responsive to verbal output (Stroop and Number) only or keypress (Flanker) only. These analyses are summarized in **Tables S5** and **S6**.

To assess the correlation between conflict responses and reaction times (**Figure 7**), a linear regression (“fitlm” function in Matlab) was performed between reaction time and the mean high-gamma or theta band power of each trial for all conflict-modulated electrodes. An electrode was considered to show a significant correlation if the p value of the linear regression slope was smaller than 0.05.

### Classifier analyses

We quantified whether we could distinguish between congruent and incongruent trials in individual trials based on the activity of pseudo-populations formed by multiple electrodes (Liu et al., 2009). We used a linear-kernel support vector machine with ten-fold cross-validation for all the classifier analyses (**Figures 8-9**). Two features were calculated for each trial from each electrode: the mean and maximum band power from average stimulus onset to the behavioral response. These analyses were conducted separately for the high-gamma and theta frequency bands. All data were normalized to zero mean and standard deviation 1 before each training and testing session. All the classifier performance results reported are based on cross-validated test data. The main text and **Figures 8-9** describe all the different combinations of training and test data used, which are critical to evaluate within-task and between-task invariance.

## Data and code availability

All the data and source code will be made available through public repositories upon publication.

## Author contributions

The task was designed by YX and GK. All the data were collected by YX with the help of CCC, NEC, IR, YCS, and DW. CRG, SS, JRM, HYY, and WSA performed the surgeries on the patients. All the data were analyzed by YX, with frequent discussions with GK. The manuscript was written by YX and GK and approved by all authors.

## Acknowledgments

This work was supported by NIH grant R01EY026025 and by the Center for Brains, Minds and Machines, funded by NSF Science and Technology Center Award CCF-1231216.

---

**Figure S1.**
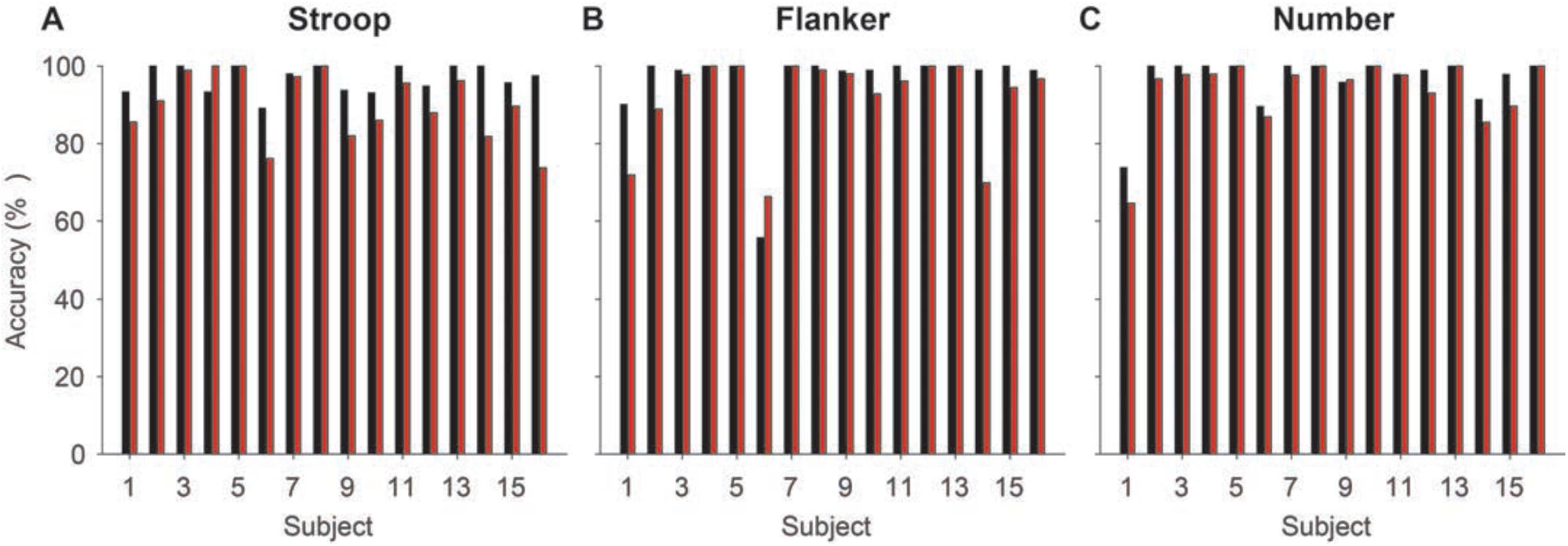
Accuracy of each subject in each task. Black bars indicate congruent trials and red bars incongruent trials.

**Figure S2.**
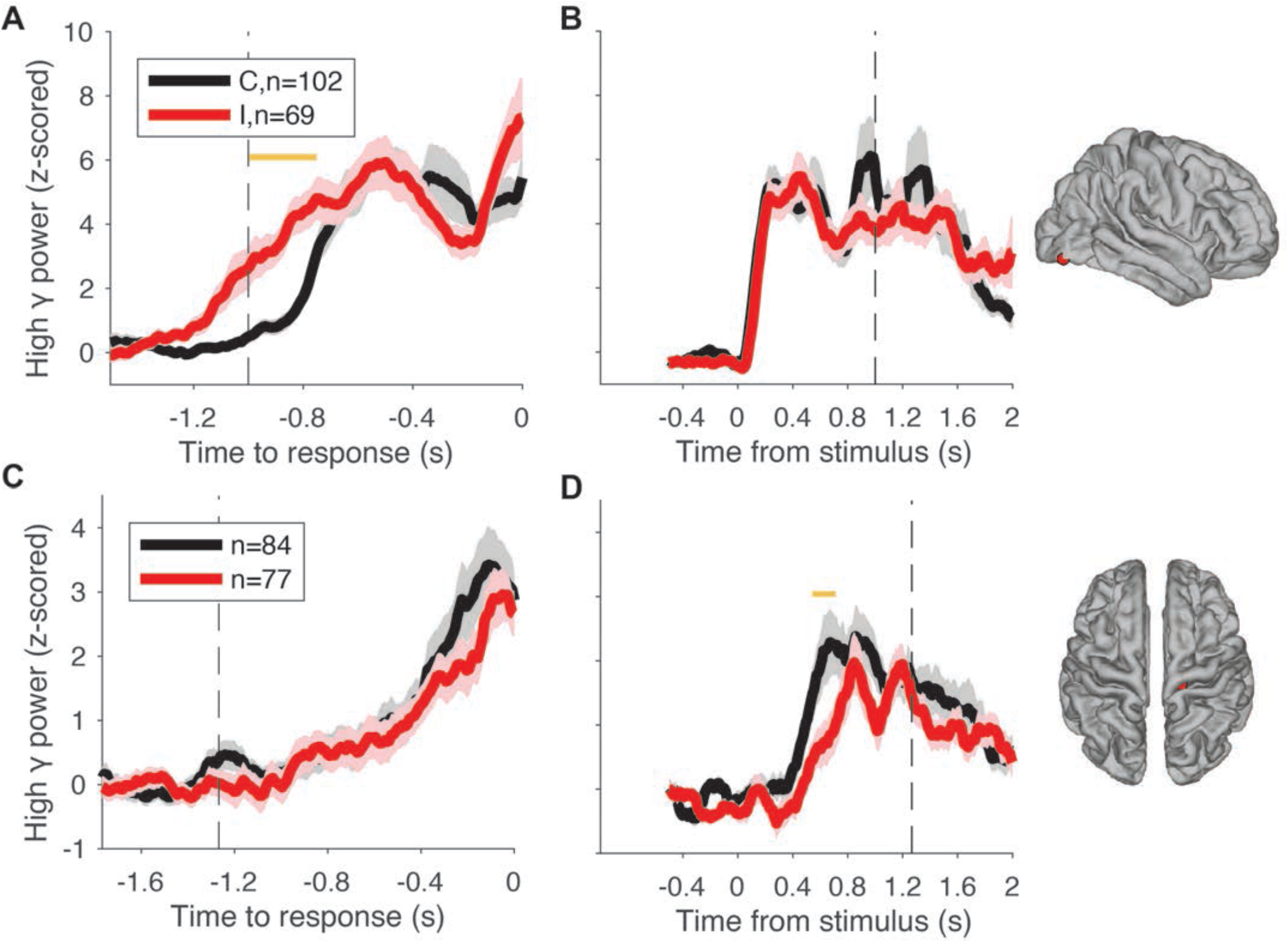
Alignment to stimulus and behavioral responses is critical to interpret conflict modulation signals. An electrode at right lateral occipital cortex showed conflict modulation when the high gamma power (mean±SEM, black for congruent and red for incongruent) was aligned to behavioral response (A) but no such effect emerged when aligned to stimulus onset (B). Conversely, an electrode at the right precentral gyrus showed conflict modulation when the high gamma power (mean±SEM) was aligned to stimulus onset (D) but not behavioral response (C). These electrodes reflect either purely visual response (B) or purely motor response (C) and thus we do not consider them as conflict-selective.

**Figure S3.**
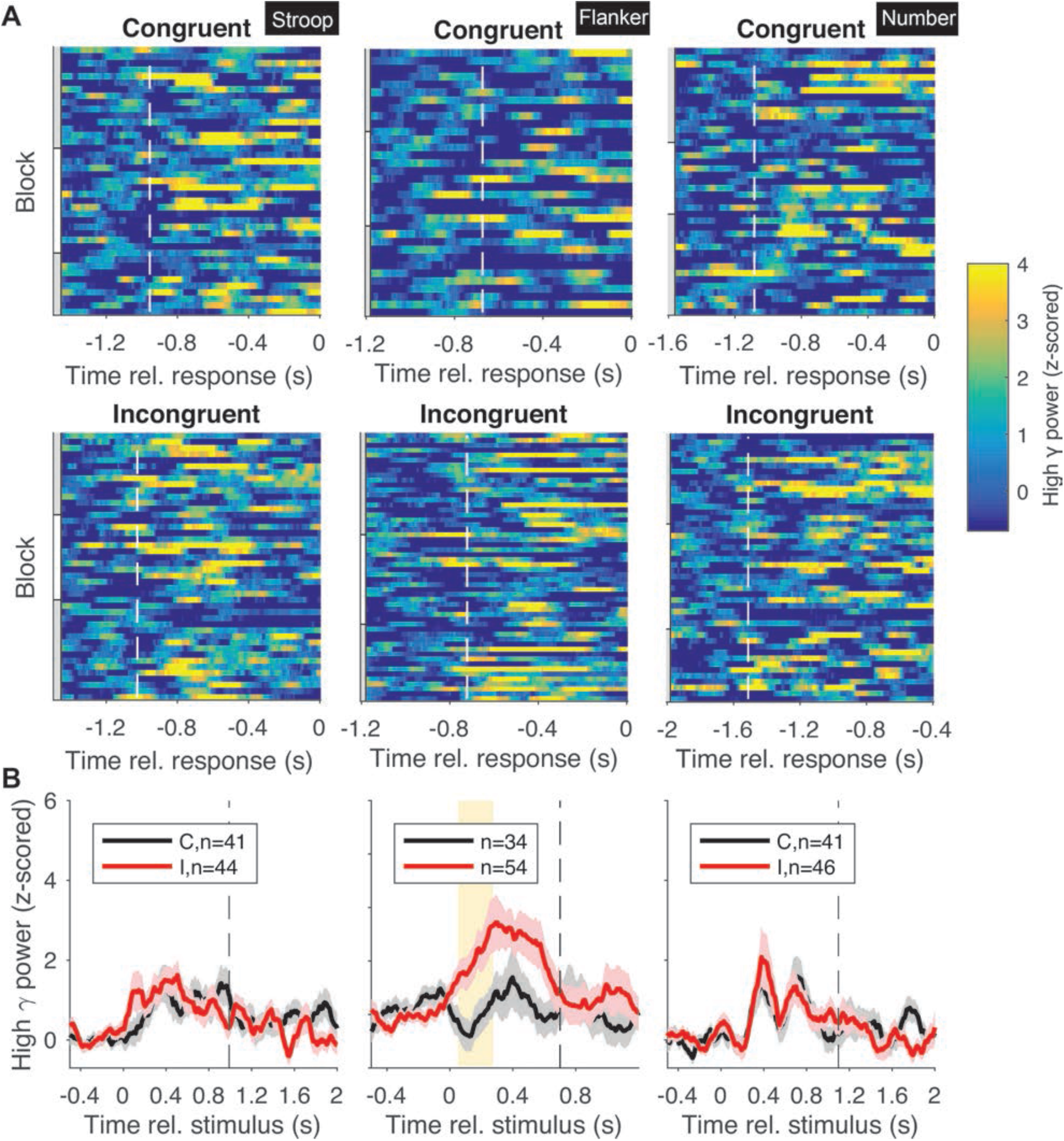
Example Flanker-specific electrode in the high gamma band (same electrode as in Figure 5A). An electrode located at the right superior parietal cortex exhibited conflict modulation in the Flanker task only. **A**. Raster plots showing the neural signals in individual trials (see color scale on the right) for congruent (top row) and incongruent (middle row) trials. The white dashed lines in the raster plots show the average stimulus onset time (these lines are shifted to the left in incongruent condition compared to congruent condition, indicating longer RT for the incongruent trials). Gray and white bars on the left of the raster plots represent different blocks. **B**. Z-scored high gamma power (mean±SEM, black for congruent and red for incongruent) aligned to stimulus onset. Vertical dashed lines denote the average behavioral response time. Yellow background indicates statistically significant difference between congruent and incongruent trials (permutation test, 5000 iterations, alpha=0.05, **Methods**).

**Figure S4.**
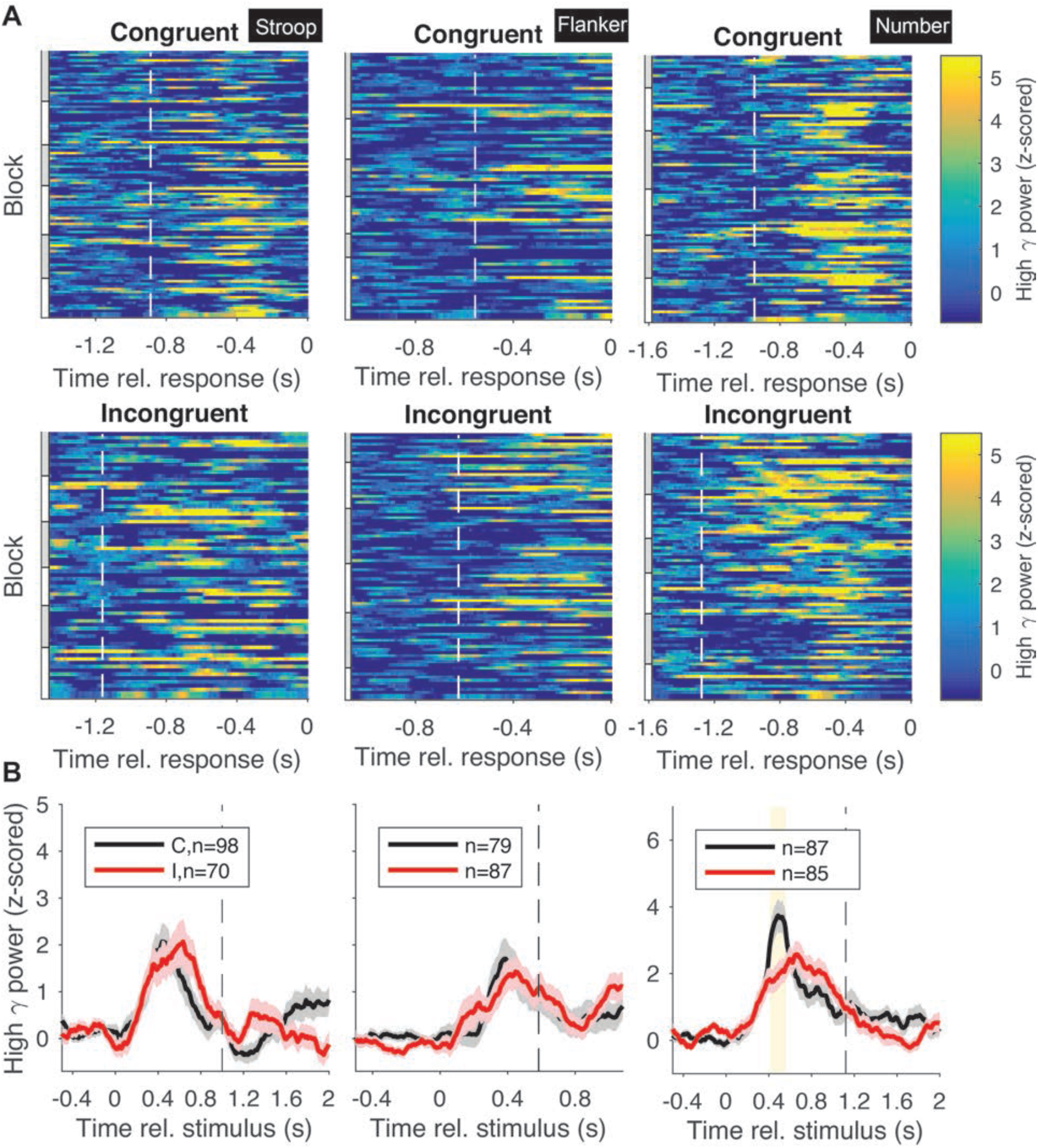
Example Number-specific electrode in the high gamma band (same as in Figure 5B). An electrode located at the right precuneus exhibited conflict modulation in the Number task only. **A**. Raster plots showing the neural signals in individual trials (see color scale on the right) for congruent (top row) and incongruent (middle row) trials. The white dashed lines in the raster plots show the average stimulus onset time (these lines are shifted to the left in incongruent condition compared to congruent condition, indicating longer RT for the incongruent trials). Gray and white bars on the left of the raster plots represent different blocks. **B**. Z-scored high gamma power (mean±SEM, black for congruent and red for incongruent) aligned to stimulus onset. Vertical dashed lines denote the average behavioral response time. Yellow background indicates statistically significant difference between congruent and incongruent trials (permutation test, 5000 iterations, alpha=0.05, **Methods**).

**Figure S5.**
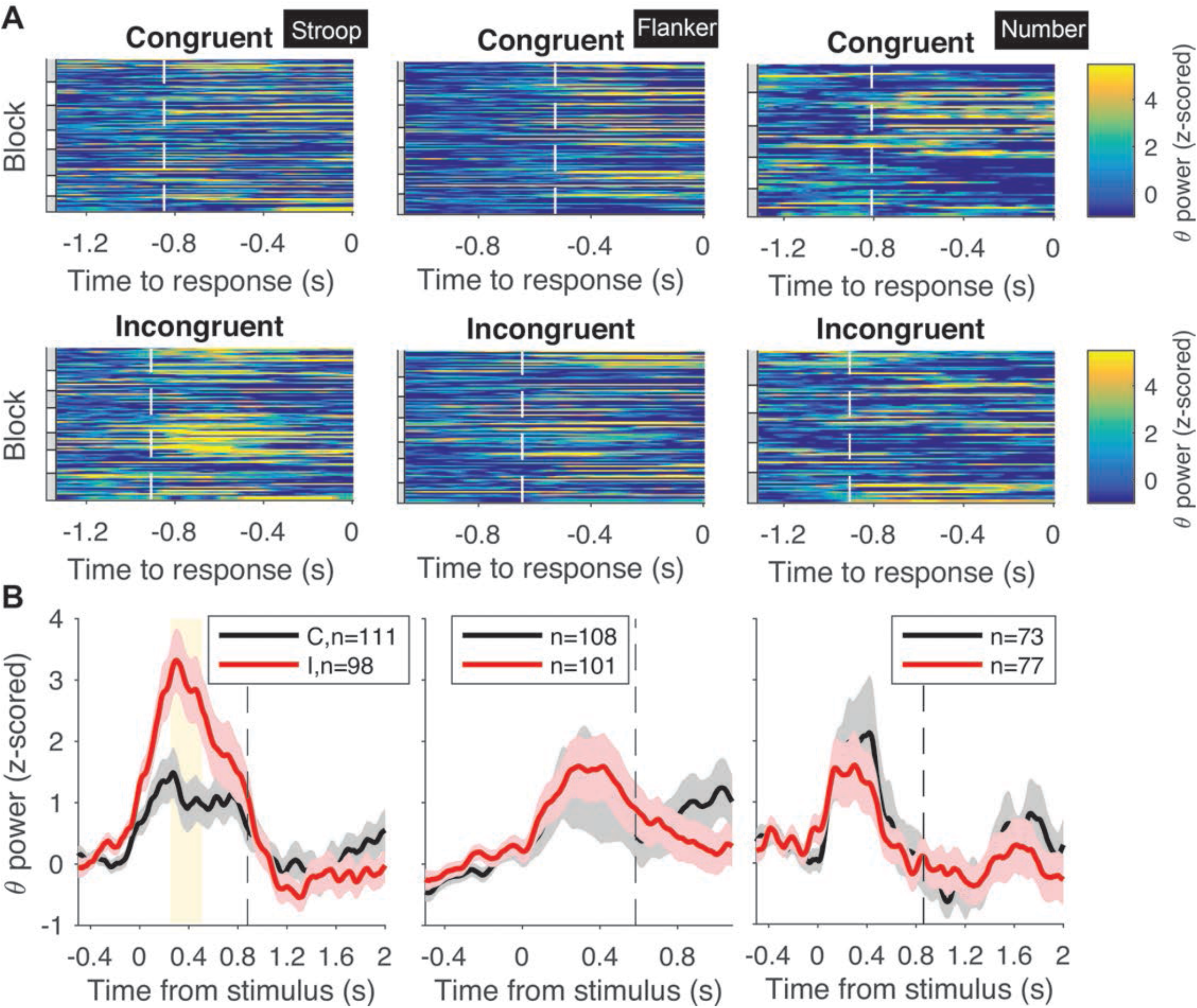
Example Stroop-specific electrode in the theta band (same as in Figure 6C). An electrode located at the right pars triangularis exhibited conflict modulation in the Stroop task only. **A**. Raster plots showing the neural signals in individual trials (see color scale on the right) for congruent (top row) and incongruent (middle row) trials. The white dashed lines in the raster plots show the average stimulus onset time (these lines are shifted to the left in incongruent condition compared to congruent condition, indicating longer RT for the incongruent trials). Gray and white bars on the left of the raster plots represent different blocks. **B**. Z-scored theta power (mean±SEM, black for congruent and red for incongruent) aligned to stimulus onset. Vertical dashed lines denote the average behavioral response time. Yellow background indicates statistically significant difference between congruent and incongruent trials (permutation test, 5000 iterations, alpha=0.05, **Methods**).

**Figure S6.**
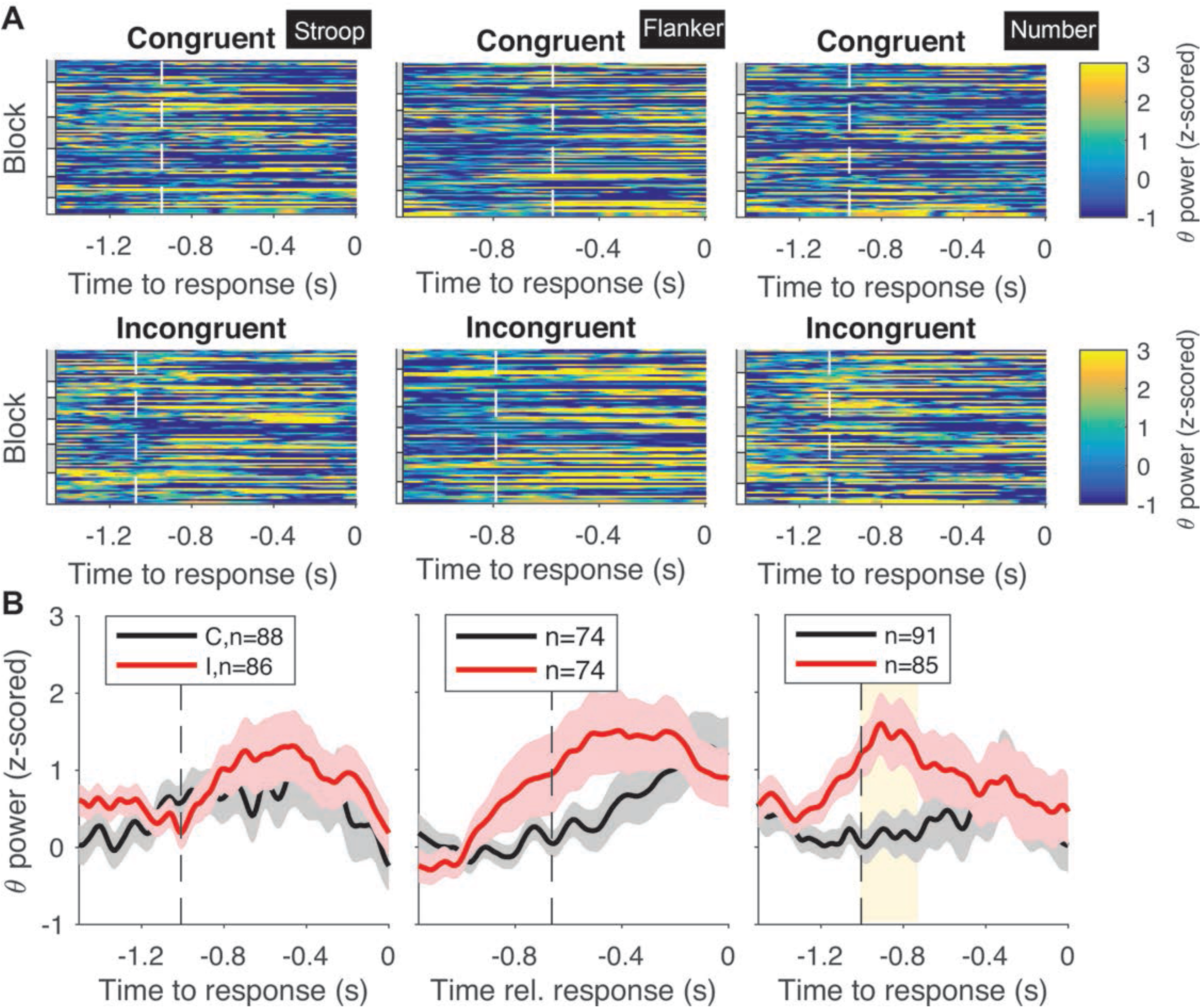
Example Number-specific electrode in the theta band (same as in Figure 6D). An electrode located at the left superior temporal cortex exhibited conflict modulation in the Number task only. **A**. Raster plots showing the neural signals in individual trials (see color scale on the right) for congruent (top row) and incongruent (middle row) trials. The white dashed lines in the raster plots show the average stimulus onset time (these lines are shifted to the left in incongruent condition compared to congruent condition, indicating longer RT for the incongruent trials). Gray and white bars on the left of the raster plots represent different blocks. **B**. Z-scored theta power (mean±SEM, black for congruent and red for incongruent) aligned to stimulus onset. Vertical dashed lines denote the average behavioral response time. Yellow background indicates statistically significant difference between congruent and incongruent trials (permutation test, 5000 iterations, alpha=0.05, **Methods**).

**Figure S7.**
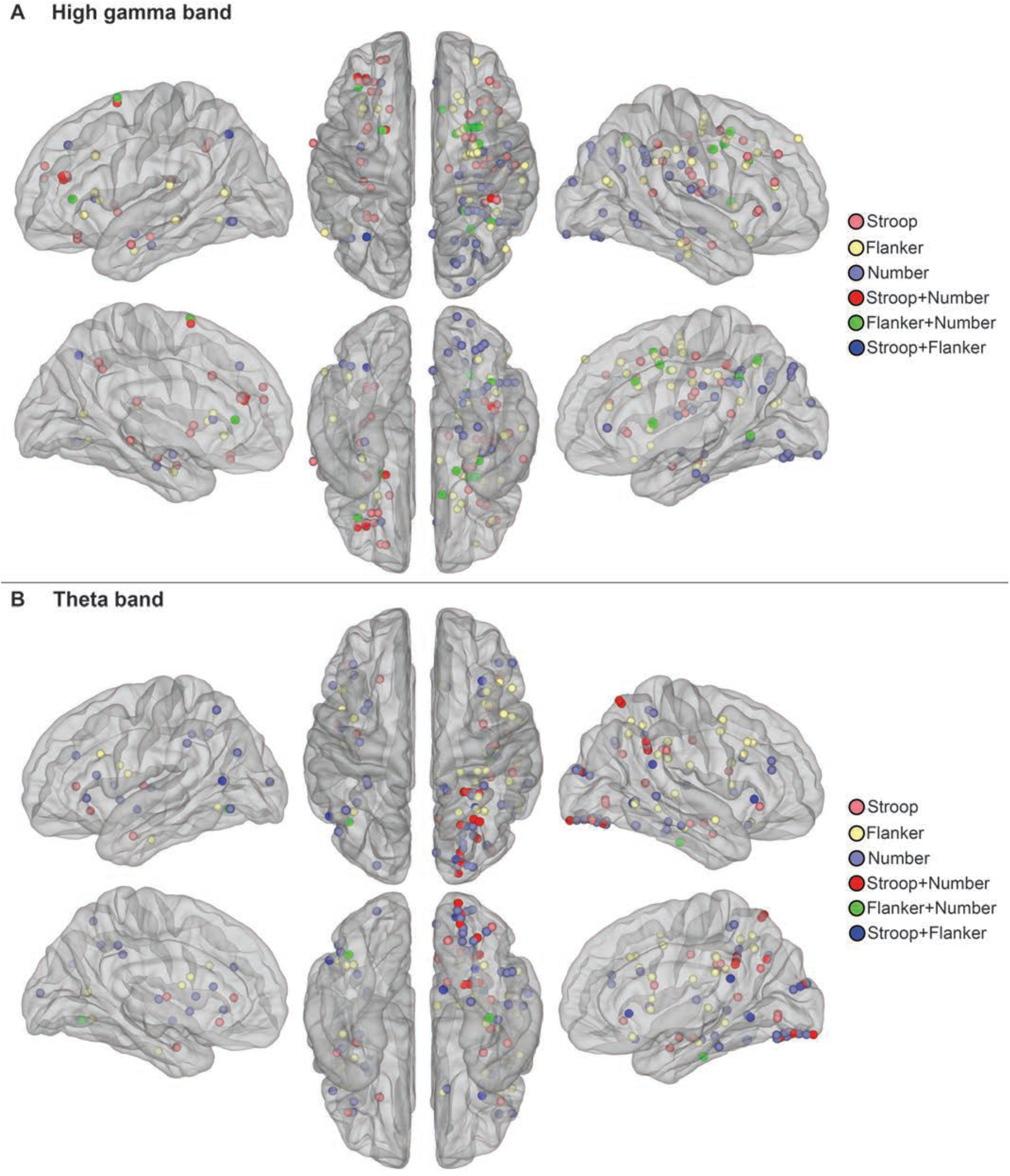
Electrodes exhibiting conflict modulation in high gamma (A) and theta (B) band. Specific locations of these electrodes can be found in Table **S3** and **S4**. We didn’t find any electrode that was conflict-selective for all the three tasks in all the frequency bands we analyzed.

**Figure S8.**
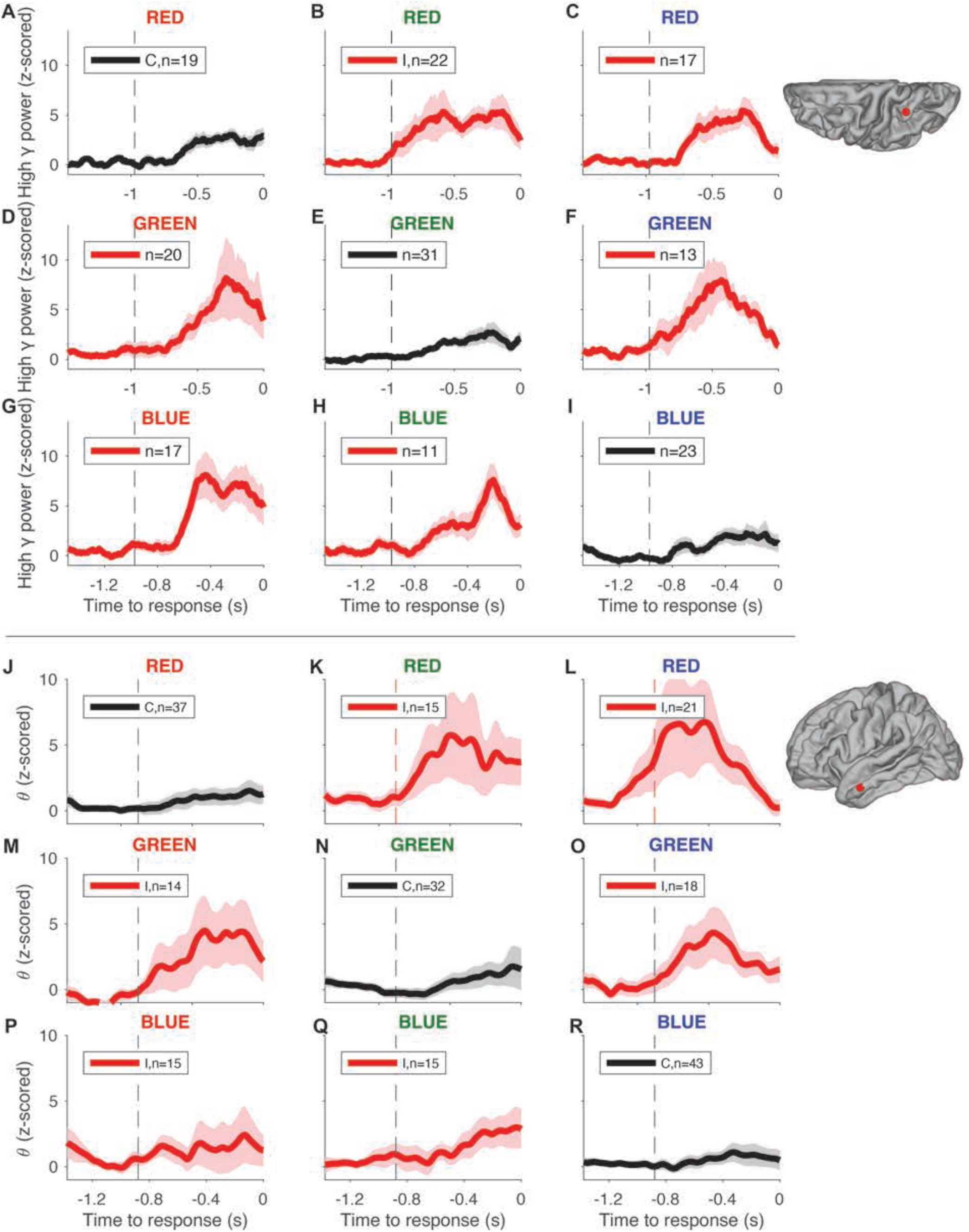
Example Stroop-specific electrodes show within-task invariance (A-I: high gamma band, left inferior parietal; J-R: theta band, left superior temporal). Z-scored high-gamma or theta power (mean±SEM) aligned to behavioral response time (black for congruent and red for incongruent). Vertical dashed lines denote the average stimulus onset. Subplot titles indicate specific stimulus types. Conflict modulation occurs in all incongruent color/word combinations.

**Figure S9.**
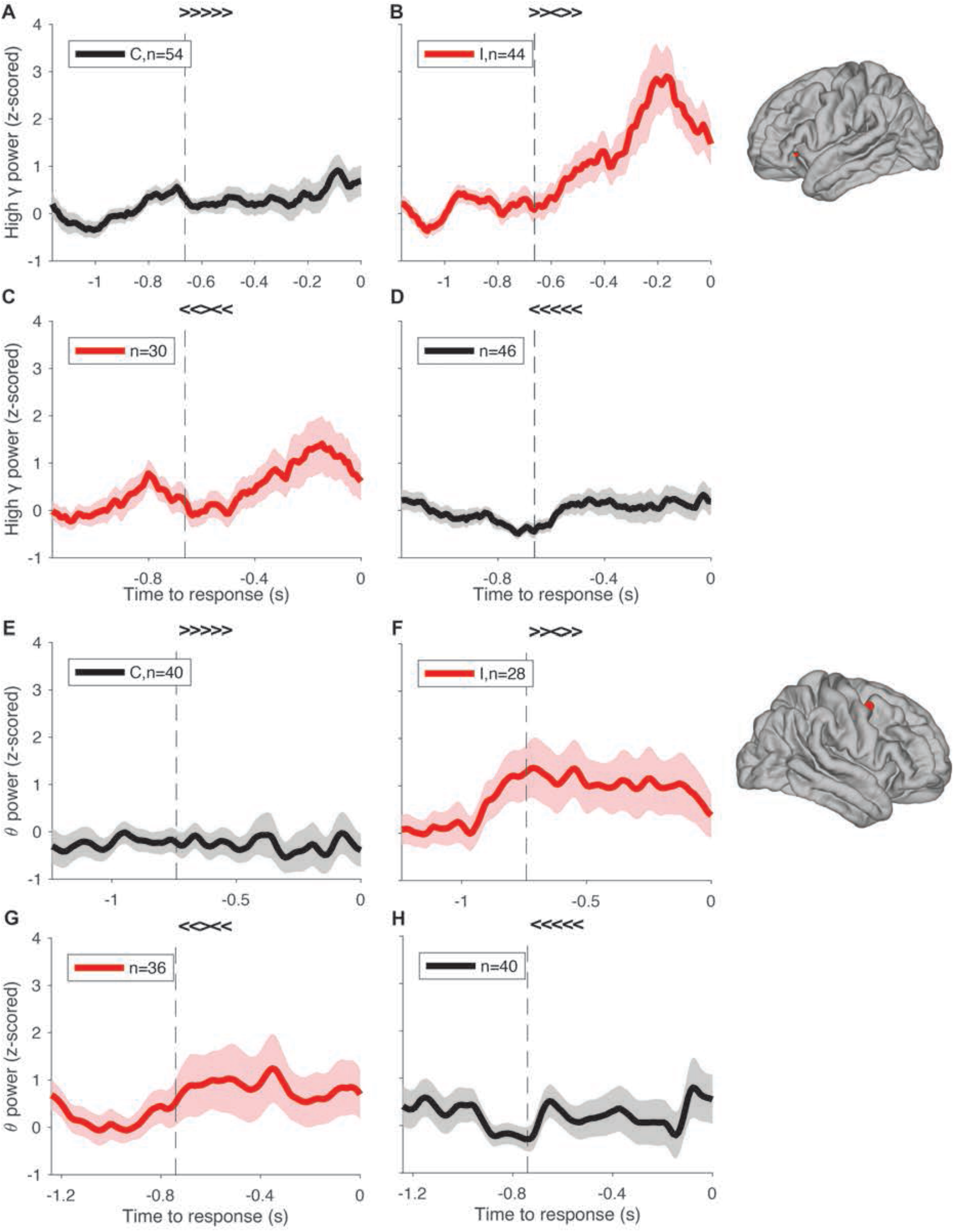
Example Flanker-specific electrodes show within-task invariance (A-D, high gamma, left orbitofrontal; E-H, theta, right precentral). Z-scored high-gamma or theta power (mean±SEM) aligned to behavioral response time (black for congruent and red for incongruent). Vertical dashed lines denote the average stimulus onset. Subplot titles indicate specific stimulus types. Conflict modulation occurs in all incongruent target/flanker combinations.

**Figure S10.**
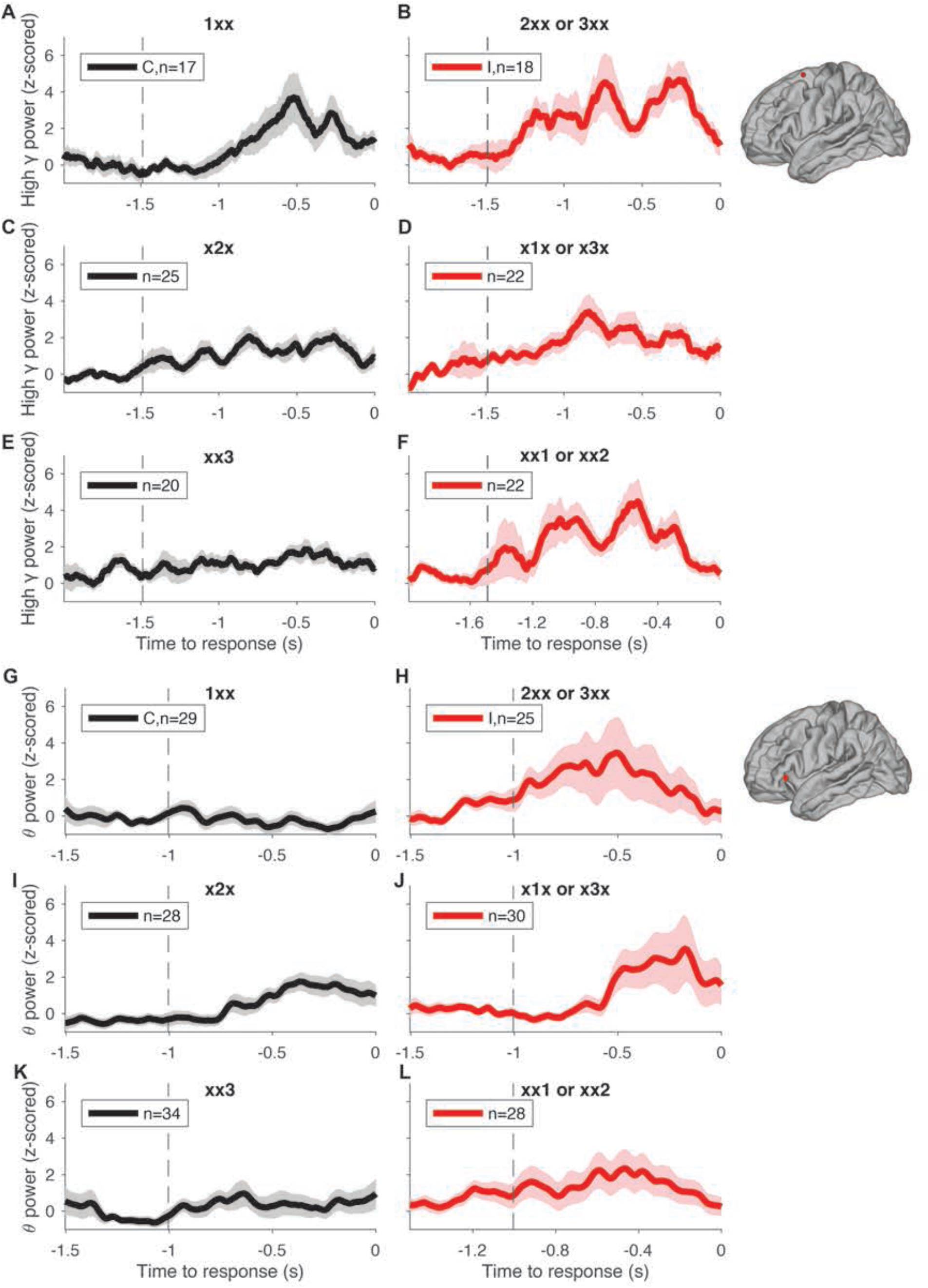
Example Number-specific electrodes show within-task invariance (A-F, high gamma, left superior frontal; G-L, theta, left pars triangularis). Z-scored high-gamma or theta power (mean±SEM) aligned to behavioral response time (black for congruent and red for incongruent). Vertical dashed lines denote the average stimulus onset. Subplot titles indicate specific stimulus types. Conflict modulation occurs in all incongruent target/distractor combinations.

**Figure S11.**
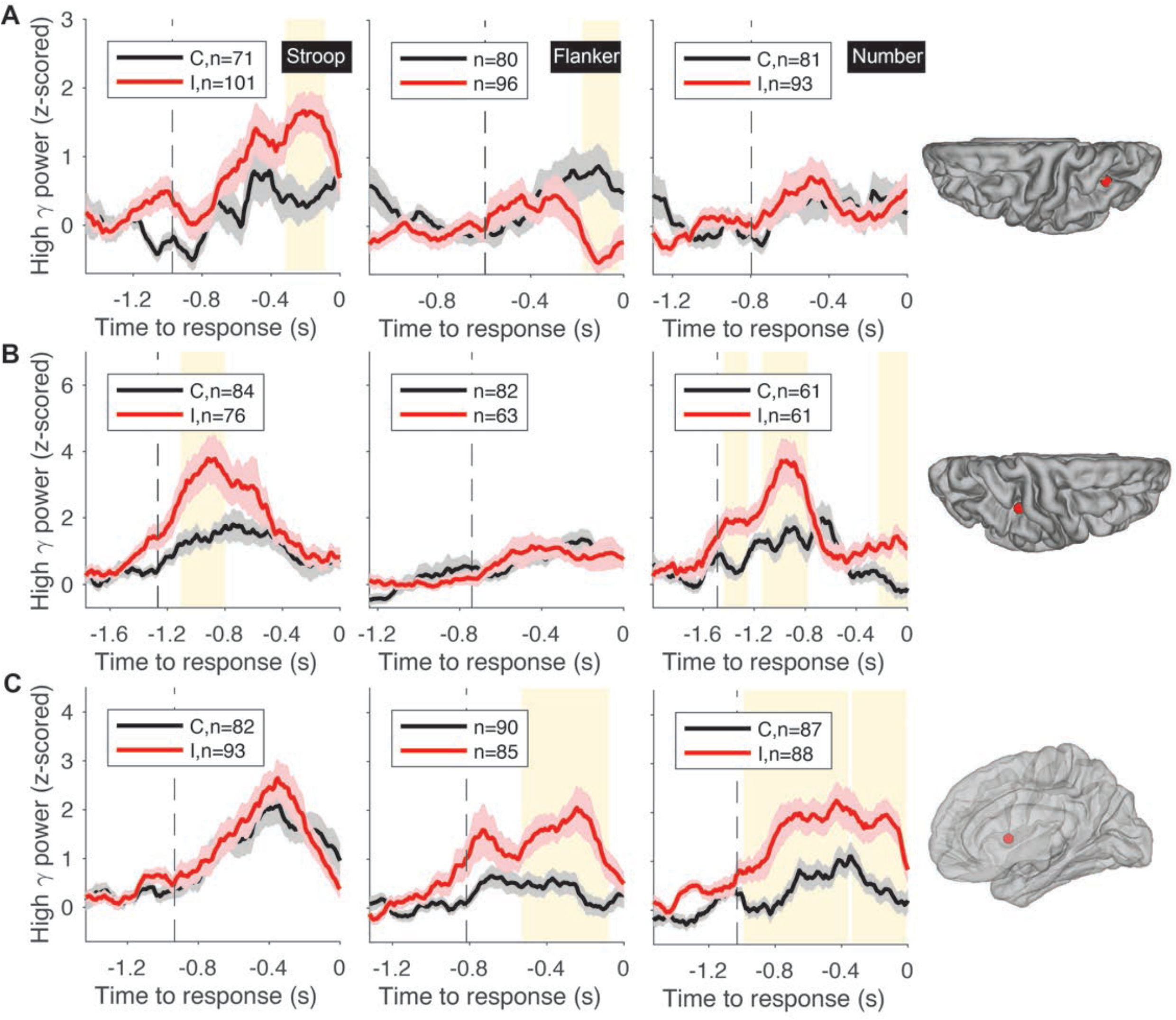
Example electrodes showing conflict modulation in two tasks. **A.** An electrode located in the left inferior parietal (see location on the right) exhibited conflict modulation in the Stroop and Flanker tasks but not in the Number task. Traces show z-scored high gamma power (mean±SEM, black for congruent and red for incongruent) aligned to behavioral response time. Vertical dashed lines denote the average stimulus onset time. Yellow background indicates statistically significant differences between congruent and incongruent trials (permutation test, 5000 iterations, alpha=0.05, **Methods**). **B**. An electrode located at the right supramarginal exhibited conflict modulation in the Stroop and Number tasks but not in the Flanker task. **C**. An electrode located in the insula exhibited conflict modulation in the Flanker and Number tasks but not in the Stroop task. Brain was rendered transparent for better visualization of electrodes in deep structures.

**Figure S12.**
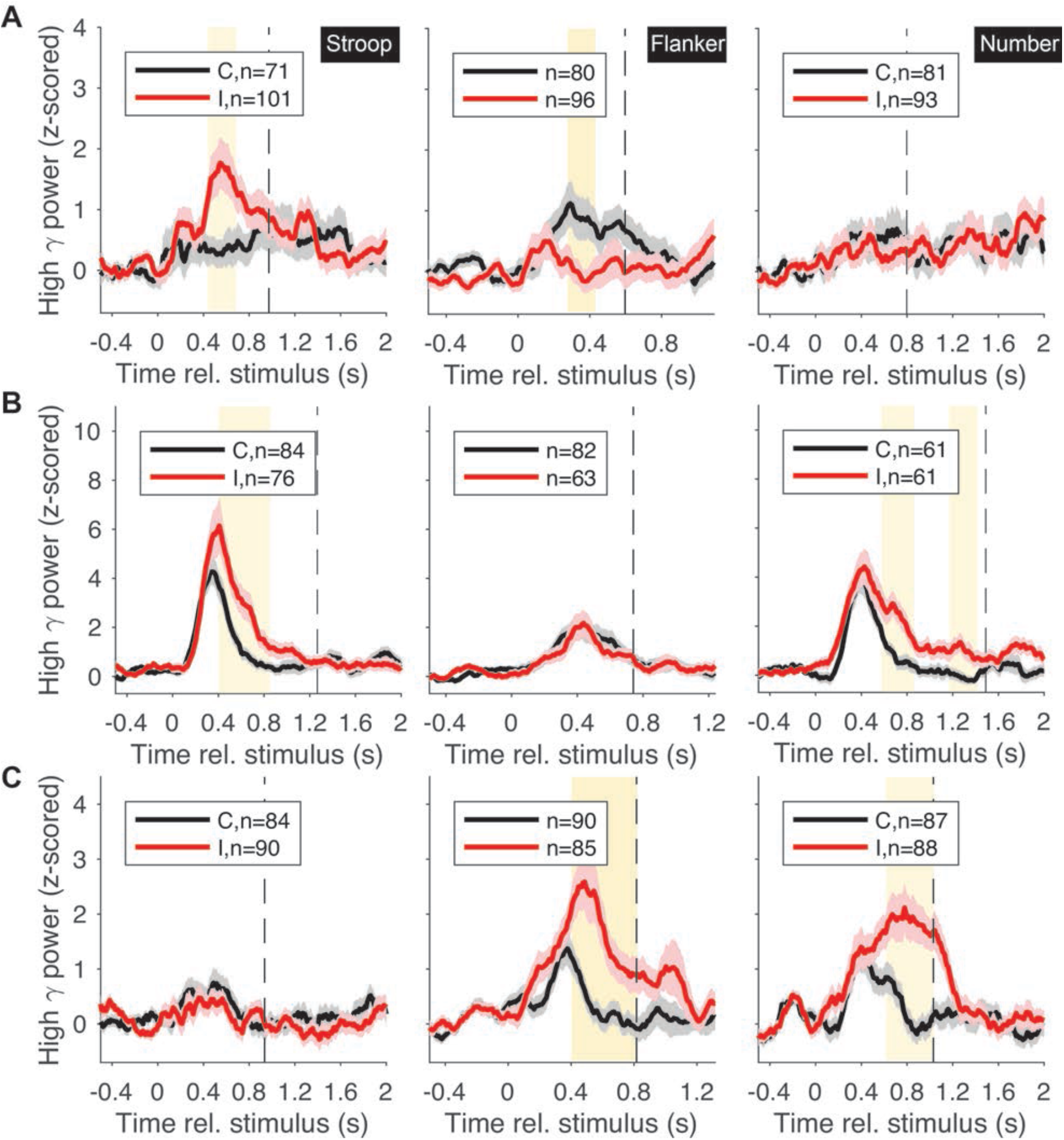
Example dual-task electrodes. Stimulus aligned responses for the electrodes in **Figure S11**.

**Figure S13.**
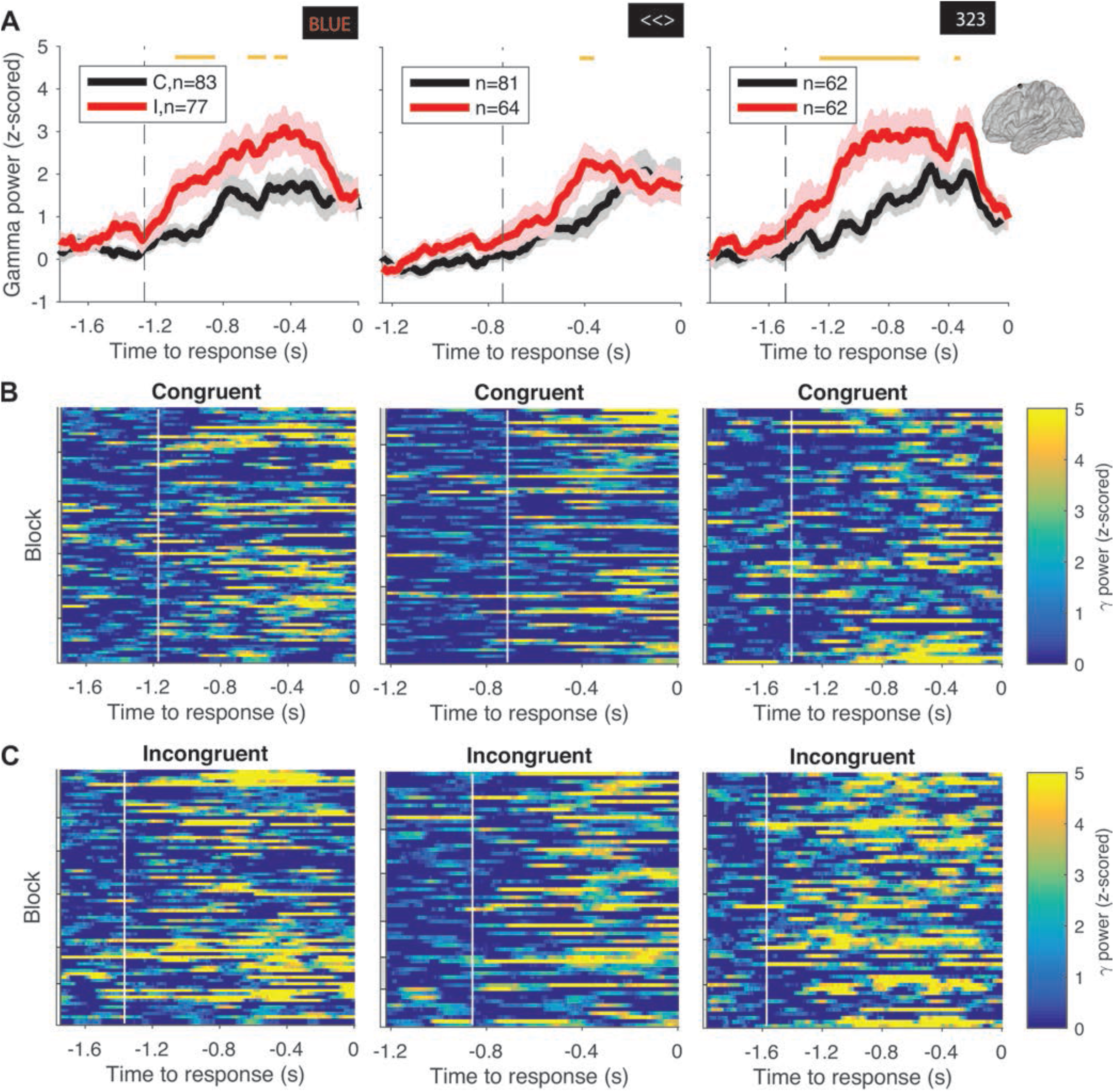
One of two task invariant electrodes. Using global referencing instead of bipolar referencing, a shorter duration threshold, and less stringent test (t-test) to distinguish incongruent and congruent trials (50 ms instead of 150 ms), we found two electrodes out of 748 electrodes (0.3%) that showed task invariance, one of them is shown here. Yellow bars indicate significant windows. This electrode was located in the left superior frontal region.

**Table S1:** Subject information. Information about each participant and number of blocks completed. Bolded entries indicate subjects that performed fewer or more than the default target number of blocks (6 blocks for each task).

| Subject Number | Age | Gender | Hospital | Number of blocks (Stroop, Flanker, Number) | Number of electrodes |
| --- | --- | --- | --- | --- | --- |
| 1 | 13 | male | BCH | 6, 6, 6 | 145 |
| 2 | 12 | female | BCH | 6, 6, 6 | 168 |
| 3 | 14 | female | BCH | 6, 6, 6 | 194 |
| 4 | 13 | male | BCH | **3, 3, 3** | 215 |
| 5 | 41 | female | BWH | **7, 7, 5** | 72 |
| 6 | 58 | female | BWH | 6, 6, 6 | 119 |
| 7 | 62 | male | JHMH | **7, 6, 6** | 99 |
| 8 | 41 | male | JHMH | 6, 6, 6 | 48 |
| 9 | 26 | male | TVGH | 6, 6, 6 | 92 |
| 10 | 27 | female | TVGH | 6, 6, 6 | 104 |
| 11 | 29 | female | TVGH | 6, 6, 6 | 101 |
| 12 | 29 | male | TVGH | 6, 6, 6 | 126 |
| 13 | 25 | male | TVGH | 6, 6, 6 | 102 |
| 14 | 20 | female | TVGH | **6, 6, 4** | 98 |
| 15 | 12 | female | TVGH | 6, 6, 6 | 90 |
| 16 | 24 | male | TVGH | 6, 6, 6 | 104 |

**Table S2:** Distribution of electrode locations. Number of electrodes in the right and left hemisphere for each location.

| **Location** | **Count** |
| --- | --- |
| 'Left-Amygdala' | 16 |
| 'Left-Hippocampus' | 16 |
| 'Left-Putamen' | 9 |
| 'Right-Amygdala' | 12 |
| 'Right-Hippocampus' | 11 |
| 'ctx-lh-bankssts' | 2 |
| 'ctx-lh-caudalanteriorcingulate' | 2 |
| 'ctx-lh-caudalmiddlefrontal' | 1 |
| 'ctx-lh-fusiform' | 5 |
| 'ctx-lh-inferiorparietal' | 22 |
| 'ctx-lh-inferiortemporal' | 10 |
| 'ctx-lh-insula' | 14 |
| 'ctx-lh-lateraloccipital' | 1 |
| 'ctx-lh-lateralorbitofrontal' | 14 |
| 'ctx-lh-lingual' | 3 |
| 'ctx-lh-medialorbitofrontal' | 6 |
| 'ctx-lh-middletemporal' | 15 |
| 'ctx-lh-parahippocampal' | 4 |
| 'ctx-lh-parsopercularis' | 3 |
| 'ctx-lh-parsorbitalis' | 1 |
| 'ctx-lh-parstriangularis' | 8 |
| 'ctx-lh-postcentral' | 2 |
| 'ctx-lh-posteriorcingulate' | 2 |
| 'ctx-lh-precentral' | 1 |
| 'ctx-lh-rostralanteriorcingulate' | 3 |
| 'ctx-lh-rostralmiddlefrontal' | 30 |
| 'ctx-lh-superiorfrontal' | 15 |
| 'ctx-lh-superiorparietal' | 4 |
| 'ctx-lh-superiortemporal' | 14 |
| 'ctx-lh-supramarginal' | 6 |
| 'ctx-rh-bankssts' | 5 |
| 'ctx-rh-caudalanteriorcingulate' | 3 |
| 'ctx-rh-caudalmiddlefrontal' | 22 |
| 'ctx-rh-cuneus' | 3 |
| 'ctx-rh-entorhinal' | 1 |
| 'ctx-rh-frontalpole' | 1 |
| 'ctx-rh-fusiform' | 19 |
| 'ctx-rh-inferiorparietal' | 21 |
| 'ctx-rh-inferiortemporal' | 18 |
| 'ctx-rh-insula' | 27 |
| 'ctx-rh-isthmuscingulate' | 8 |
| 'ctx-rh-lateraloccipital' | 7 |
| 'ctx-rh-lateralorbitofrontal' | 11 |
| 'ctx-rh-lingual' | 7 |
| 'ctx-rh-medialorbitofrontal' | 5 |
| 'ctx-rh-middletemporal' | 23 |
| 'ctx-rh-paracentral' | 9 |
| 'ctx-rh-parahippocampal' | 6 |
| 'ctx-rh-parsopercularis' | 9 |
| 'ctx-rh-parsorbitalis' | 3 |
| 'ctx-rh-parstriangularis' | 5 |
| 'ctx-rh-pericalcarine' | 6 |
| 'ctx-rh-postcentral' | 29 |
| 'ctx-rh-posteriorcingulate' | 8 |
| 'ctx-rh-precentral' | 43 |
| 'ctx-rh-precuneus' | 13 |
| 'ctx-rh-rostralanteriorcingulate' | 3 |
| 'ctx-rh-rostralmiddlefrontal' | 15 |
| 'ctx-rh-superiorfrontal' | 26 |
| 'ctx-rh-superiorparietal' | 33 |
| 'ctx-rh-superiortemporal' | 21 |
| 'ctx-rh-supramarginal' | 29 |
| 'ctx-rh-temporalpole' | 1 |
| 'ctx-rh-transversetemporal' | 2 |
| **Total** | **694** |

**Table S3:** Location and specificity of conflict-modulated electrodes (high-gamma band). For each location, the table reports the number of electrodes that show conflict modulation in one task only, in two tasks, or in all three tasks. Sum refers to the total number of conflict-modulated electrodes in each row and ‘total in location’ shows the total number of electrodes analyzed in that regions. S=Stroop, F=Flanker, N=Number.

| Location | One task only |  |  | Two tasks |  |  | All | Sum |
| --- | --- | --- | --- | --- | --- | --- | --- | --- |
|  | Stroop | Flanker | Number | S+F | S+N | F+N | S+F+N |  |
| Amygdala |  | 1 | 1 |  |  |  |  | 2 |
| ctx-caudalmiddlefrontal | 1 | 1 |  |  |  |  |  | 2 |
| ctx-entorhinal |  |  | 1 |  |  |  |  | 1 |
| ctx-fusiform |  | 2 | 4 |  |  |  |  | 6 |
| ctx-inferiorparietal | 3 | 5 | 2 | 1 |  |  |  | 11 |
| ctx-inferiortemporal |  | 2 | 6 |  |  | 1 |  | 9 |
| ctx-insula | 1 | 3 | 3 |  |  | 2 |  | 9 |
| ctx-lateraloccipital |  |  | 3 |  |  |  |  | 3 |
| ctx-lateralorbitofrontal | 4 | 2 |  |  |  |  |  | 6 |
| ctx-midtemporal | 2 | 1 | 1 |  |  |  |  | 4 |
| ctx-paracentral | 1 |  |  |  |  |  |  | 1 |
| ctx-parahippocampal | 1 |  | 1 |  |  |  |  | 2 |
| ctx-parstriangularis |  | 1 |  |  |  | 1 |  | 2 |
| ctx-pericalcarine |  |  | 1 |  |  |  |  | 1 |
| ctx-postcentral | 2 |  | 2 |  |  |  |  | 4 |
| ctx-posteriorcingulate |  | 1 | 1 |  |  |  |  | 2 |
| ctx-precentral | 4 | 5 | 1 |  |  | 1 |  | 11 |
| ctx-precuneus |  |  | 2 |  |  |  |  | 2 |
| ctx- rostralanteriorcingulate |  |  | 1 |  |  |  |  | 1 |
| ctx-rostralmiddlefrontal | 6 | 2 |  |  | 2 |  |  | 10 |
| ctx-superiorfrontal | 3 | 5 | 1 |  | 1 | 4 |  | 14 |
| ctx-superiorparietal | 1 | 2 | 10 |  |  | 2 |  | 15 |
| ctx-superiortemporal | 1 | 1 |  |  |  |  |  | 2 |
| ctx-supramarginal | 4 | 3 | 2 |  | 1 |  |  | 10 |
| Hippocampus | 1 |  | 1 |  |  |  |  | 2 |
| Putamen | 2 |  |  |  |  |  |  | 2 |
| **Total** | **37** | **37** | **44** | **1** | **4** | **11** | **0** | **134** |
|  | **118** |  |  | **16** |  |  | **0** | **134** |

**Table S4:** Location and specificity of conflict-modulated electrodes (theta band). For each location, the table reports the number of electrodes that show conflict modulation in one task only, in two tasks, or in all three tasks. Sum refers to the total number of conflict-modulated electrodes in each row and ‘total in location’ shows the total number of electrodes analyzed in that regions. S=Stroop, F=Flanker, N=Number.

| Location | One task only |  |  | Two tasks |  |  | All | Sum |
| --- | --- | --- | --- | --- | --- | --- | --- | --- |
|  | Stroop | Flanker | Number | S+F | S+N | F+N | S+F+N |  |
| Amygdala | 1 |  |  |  |  |  |  | 1 |
| ctx-bankssts |  |  | 1 |  |  |  |  | 1 |
| ctx-caudalmiddlefrontal | 1 | 3 | 1 |  |  |  |  | 5 |
| ctx-cuneus |  |  | 1 |  |  |  |  | 1 |
| ctx-fusiform | 1 | 1 | 5 |  | 1 | 1 |  | 9 |
| ctx-inferiorparietal |  | 3 | 3 | 1 |  |  |  | 7 |
| ctx-inferiortemporal |  | 2 | 5 |  |  | 1 |  | 8 |
| ctx-insula | 1 | 2 | 1 |  |  |  |  | 4 |
| ctx-isthmuscingulate | 1 | 1 |  | 1 |  |  |  | 3 |
| ctx-lateraloccipital |  |  | 4 | 1 | 2 |  |  | 7 |
| ctx-lateralorbitofrontal | 1 |  |  | 1 |  |  |  | 2 |
| ctx-lingual |  | 1 | 2 | 1 | 1 |  |  | 5 |
| ctx-medialorbitofrontal |  | 1 | 1 |  |  |  |  | 2 |
| ctx-middletemporal |  | 1 | 1 |  |  |  |  | 2 |
| ctx-parahippocampal |  | 1 | 1 |  |  |  |  | 2 |
| ctx-parsopercularis |  | 1 | 2 |  |  |  |  | 3 |
| ctx-parstriangularis | 2 |  | 1 |  |  |  |  | 3 |
| ctx-pericalcarine | 2 |  |  |  |  |  |  | 2 |
| ctx-postcentral | 1 | 1 |  |  |  |  |  | 2 |
| ctx-precentral |  | 3 |  |  |  |  |  | 3 |
| ctx-precuneus | 1 |  | 1 |  | 1 |  |  | 3 |
| ctx-rostralmiddlefrontal |  | 2 | 3 |  |  |  |  | 5 |
| ctx-superiorparietal |  | 4 | 7 |  | 5 |  |  | 16 |
| ctx-superiortemporal | 2 | 1 | 1 |  |  |  |  | 4 |
| ctx-supramarginal | 2 | 2 |  |  |  |  |  | 4 |
| Hippocampus | 1 | 1 |  |  |  |  |  | 2 |
| Putamen |  |  | 3 |  |  |  |  | 3 |
| **Total** | **17** | **31** | **44** | **5** | **10** | **2** | **0** | **109** |
|  | **92** |  |  | **17** |  |  | **0** | **109** |

**Table S5:**
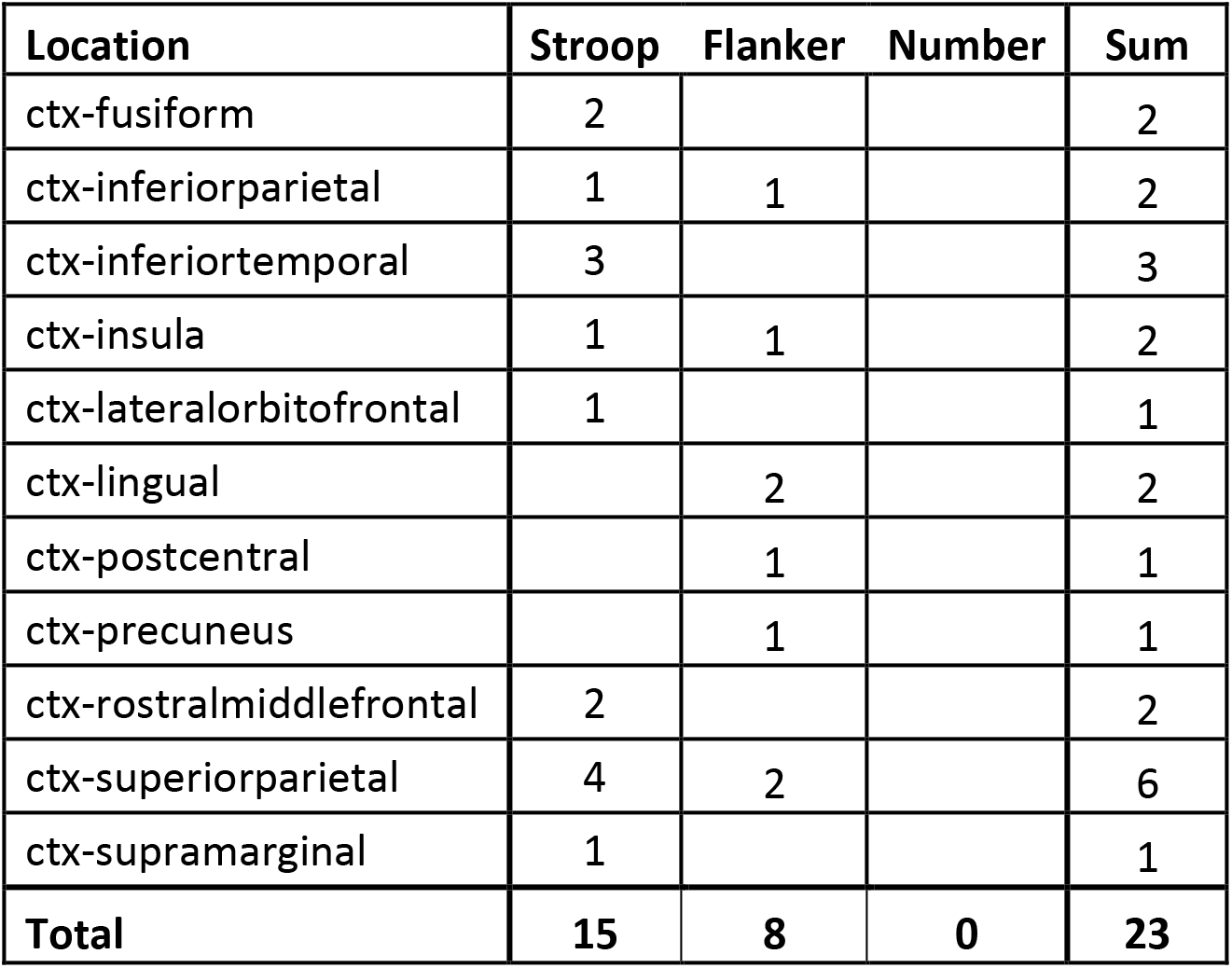
Location of visual-selective electrodes. For each location, the table reports the number of electrodes that show visual selectivity during Stroop, Flanker, and Number task respectively. Sum refers to the total number of response electrodes in each row and ‘total in location’ shows the total number of electrodes analyzed in that region.

| Location | Stroop | Flanker | Number | Sum |
| --- | --- | --- | --- | --- |
| ctx-fusiform | 2 |  |  | 2 |
| ctx-inferiorparietal | 1 | 1 |  | 2 |
| ctx-inferiortemporal | 3 |  |  | 3 |
| ctx-insula | 1 | 1 |  | 2 |
| ctx-lateralorbitofrontal | 1 |  |  | 1 |
| ctx-lingual |  | 2 |  | 2 |
| ctx-postcentral |  | 1 |  | 1 |
| ctx-precuneus |  | 1 |  | 1 |
| ctx-rostralmiddlefrontal | 2 |  |  | 2 |
| ctx-superiorparietal | 4 | 2 |  | 6 |
| ctx-supramarginal | 1 |  |  | 1 |
| **Total** | **15** | **8** | **0** | **23** |

**Table S6:** Location of motor-selective electrodes. For each location, the table reports the number of electrodes that show motor selectivity for verbal and keypress. Sum refers to the total number of response electrodes in each row and ‘total in location’ shows the total number of electrodes analyzed in that region.

| Location | Verbal | Keypress | Sum |
| --- | --- | --- | --- |
| ctx-caudalanteriorcingulate |  | 1 | 1 |
| ctx-caudalmiddlefrontal | 1 |  | 1 |
| ctx-inferiorparietal |  | 2 | 2 |
| ctx-insula | 1 |  | 1 |
| ctx-parahippocampal |  | 1 | 1 |
| ctx-parsopercularis | 1 |  | 1 |
| ctx-parstriangularis | 1 |  | 1 |
| ctx-precentral | 7 | 1 | 8 |
| ctx-postcentral | 11 | 2 | 13 |
| ctx-superiorfrontal | 1 |  | 1 |
| ctx-superiorparietal |  | 1 | 1 |
| ctx-superiortemporal | 2 |  | 2 |
| ctx-supramarginal | 1 | 2 | 3 |
| **Total** | **26** | **10** | **36** |

**Table S7:** umber of conflict-modulated electrodes considering other frequency bands.

| Frequency Band | Time-bandwidth product, taper, moving window | One task only |  |  | Two tasks |  |  | All | Sum |
| --- | --- | --- | --- | --- | --- | --- | --- | --- | --- |
|  |  | Stroop | Flanker | Number | S+F | S+N | F+N | S+F+N |  |
| High-gamma (70-120 Hz) | 5, 7, 200 ms | 37 | 37 | 44 | 1 | 4 | 11 | 0 | 134 |
| Low gamma (35-70 Hz) | 5, 7, 200 ms | 11 | 30 | 14 | 4 | 0 | 0 | 0 | 59 |
| Beta (12-35 Hz) | 3, 5, 200 ms | 13 | 26 | 39 | 0 | 4 | 1 | 0 | 83 |
| Alpha (8-12 Hz) | 2, 3, 500 ms | 16 | 40 | 23 | 1 | 3 | 7 | 0 | 90 |
| Theta (4-8 Hz) | 2, 3, 500 ms | 17 | 31 | 44 | 5 | 10 | 2 | 0 | 109 |

## References

Bansal, A.K., Singer, J.M., Anderson, W.S., Golby, A., Madsen, J.R., and Kreiman, G. (2012). Temporal stability of visually selective responses in intracranial field potentials recorded from human occipital and temporal lobes. J Neurophysiol 108, 3073–3086.

Blackman, R.K., Crowe, D.A., DeNicola, A.L., Sakellaridi, S., MacDonald, A.W., 3rd, and Chafee, M.V. (2016). Monkey Prefrontal Neurons Reflect Logical Operations for Cognitive Control in a Variant of the AX Continuous Performance Task (AX-CPT). J Neurosci 36, 4067–4079.

Botvinick, M.M., Braver, T.S., Barch, D.M., Carter, C.S., and Cohen, J.D. (2001). Conflict monitoring and cognitive control. Psychol Rev 108, 624–652.

Bunge, S.A., Hazeltine, E., Scanlon, M.D., Rosen, A.C., and Gabrieli, J.D. (2002). Dissociable contributions of prefrontal and parietal cortices to response selection. Neuroimage 17, 1562–1571.

Bush, G., and Shin, L.M. (2006). The Multi-Source Interference Task: an fMRI task that reliably activates the cingulo-frontal-parietal cognitive/attention network. Nature protocols 1, 308–313.

Carter, C.S., and van Veen, V. (2007). Anterior cingulate cortex and conflict detection: an update of theory and data. Cogn Affect Behav Neurosci 7, 367–379.

Caruana, F., Uithol, S., Cantalupo, G., Sartori, I., Lo Russo, G., and Avanzini, P. (2014). How action selection can be embodied: intracranial gamma band recording shows response competition during the Eriksen flankers test. Front Hum Neurosci 8, 668.

Cavanagh, J.F., and Frank, M.J. (2014). Frontal theta as a mechanism for cognitive control. Trends Cogn Sci 18, 414–421.

Cole, M.W., Yeung, N., Freiwald, W.A., and Botvinick, M. (2009). Cingulate cortex: diverging data from humans and monkeys. Trends Neurosci 32, 566–574.

Coulthard, E.J., Nachev, P., and Husain, M. (2008). Control over conflict during movement preparation: role of posterior parietal cortex. Neuron 58, 144–157.

Crone, N.E., Miglioretti, D.L., Gordon, B., and Lesser, R.P. (1998). Functional mapping of human sensorimotor cortex with electrocorticographic spectral analysis. II. Event-related synchronization in the gamma band. Brain : a journal of neurology 121 ( *Pt 12*), 2301–2315.

Dale, A.M., Fischl, B., and Sereno, M.I. (1999). Cortical surface-based analysis. I. Segmentation and surface reconstruction. Neuroimage 9, 179–194.

Davelaar, E.J., and Stevens, J. (2009). Sequential dependencies in the Eriksen flanker task: a direct comparison of two competing accounts. Psychon Bull Rev 16, 121–126.

Desikan, R.S., Segonne, F., Fischl, B., Quinn, B.T., Dickerson, B.C., Blacker, D., Buckner, R.L., Dale, A.M., Maguire, R.P., Hyman, B.T., et al. (2006). An automated labeling system for subdividing the human cerebral cortex on MRI scans into gyral based regions of interest. Neuroimage 31, 968–980.

Diamond, A. (2013). Executive functions. Annual review of psychology 64, 135–168.

Dosenbach, N.U., Fair, D.A., Miezin, F.M., Cohen, A.L., Wenger, K.K., Dosenbach, R.A., Fox, M.D., Snyder, A.Z., Vincent, J.L., Raichle, M.E., et al. (2007). Distinct brain networks for adaptive and stable task control in humans. Proc Natl Acad Sci U S A 104, 11073–11078.

Dosenbach, N.U., Visscher, K.M., Palmer, E.D., Miezin, F.M., Wenger, K.K., Kang, H.C., Burgund, E.D., Grimes, A.L., Schlaggar, B.L., and Petersen, S.E. (2006). A core system for the implementation of task sets. Neuron 50, 799–812.

Ebitz, R.B., Smith, E.H., Horga, G., Schevon, C.A., Yates, M.J., McKhann, G.M., Botvinick, M.M., Sheth, S.A., and Hayden, B.Y. (2020). Human dorsal anterior cingulate neurons signal conflict by amplifying task-relevant information. bioRxiv, 2020.2003.2014.991745.

Egner, T., and Hirsch, J. (2005). Cognitive control mechanisms resolve conflict through cortical amplification of task-relevant information. Nat Neurosci 8, 1784–1790.

Eriksen, B.A., and Eriksen, C.W. (1974). Effects of noise letters upon the identification of a target letter. Perception & Psychophysics 16, 143–149.

Fan, J., Flombaum, J.I., McCandliss, B.D., Thomas, K.M., and Posner, M.I. (2003). Cognitive and brain consequences of conflict. Neuroimage 18, 42–57.

Fried, I., Rutishauser, U., Cerf, M., and Kreiman, G. (2014). Single neuron studies of the human brain : probing cognition (Cambridge, Massachusetts: The MIT Press).

Fu, Z., Wu, D.J., Ross, I., Chung, J.M., Mamelak, A.N., Adolphs, R., and Rutishauser, U. (2019). Single-Neuron Correlates of Error Monitoring and Post-Error Adjustments in Human Medial Frontal Cortex. Neuron 101, 165–177 e165.

Goghari, V.M., and MacDonald, A.W., 3rd (2009). The neural basis of cognitive control: response selection and inhibition. Brain Cogn 71, 72–83.

Goschke, T. (2014). Dysfunctions of decision-making and cognitive control as transdiagnostic mechanisms of mental disorders: advances, gaps, and needs in current research. Int J Methods Psychiatr Res 23 Suppl 1, 41–57.

Gratton, G., Coles, M.G., and Donchin, E. (1992). Optimizing the use of information: strategic control of activation of responses. J Exp Psychol Gen 121, 480–506.

Gratton, G., Cooper, P., Fabiani, M., Carter, C.S., and Karayanidis, F. (2018). Dynamics of cognitive control: Theoretical bases, paradigms, and a view for the future. Psychophysiology 55.

Groppe, D.M., Bickel, S., Dykstra, A.R., Wang, X., Megevand, P., Mercier, M.R., Lado, F.A., Mehta, A.D., and Honey, C.J. (2017). iELVis: An open source MATLAB toolbox for localizing and visualizing human intracranial electrode data. J Neurosci Methods 281, 40–48.

Hanslmayr, S., Pastotter, B., Bauml, K.H., Gruber, S., Wimber, M., and Klimesch, W. (2008). The electrophysiological dynamics of interference during the Stroop task. J Cogn Neurosci 20, 215–225.

Heilbronner, S.R., and Hayden, B.Y. (2016). Dorsal Anterior Cingulate Cortex: A Bottom-Up View. Annu Rev Neurosci 39, 149–170.

Helfrich, R.F., and Knight, R.T. (2016). Oscillatory Dynamics of Prefrontal Cognitive Control. Trends Cogn Sci 20, 916–930.

Janssens, C., De Loof, E., Boehler, C.N., Pourtois, G., and Verguts, T. (2018). Occipital alpha power reveals fast attentional inhibition of incongruent distractors. Psychophysiology 55.

Joshi, A., Scheinost, D., Okuda, H., Belhachemi, D., Murphy, I., Staib, L.H., and Papademetris, X. (2011). Unified framework for development, deployment and robust testing of neuroimaging algorithms. Neuroinformatics 9, 69–84.

Koga, S., Rothermel, R., Juhasz, C., Nagasawa, T., Sood, S., and Asano, E. (2011). Electrocorticographic correlates of cognitive control in a Stroop task-intracranial recording in epileptic patients. Hum Brain Mapp 32, 1580–1591.

Lesh, T.A., Niendam, T.A., Minzenberg, M.J., and Carter, C.S. (2011). Cognitive control deficits in schizophrenia: mechanisms and meaning. Neuropsychopharmacology 36, 316–338.

Li, Y.S., Nassar, M.R., Kable, J.W., and Gold, J.I. (2019). Individual Neurons in the Cingulate Cortex Encode Action Monitoring, Not Selection, during Adaptive Decision-Making. J Neurosci 39, 6668–6683.

Liu, H., Agam, Y., Madsen, J.R., and Kreiman, G. (2009). Timing, timing, timing: fast decoding of object information from intracranial field potentials in human visual cortex. Neuron 62, 281–290.

MacLeod, C.M. (1991). Half a century of research on the Stroop effect: an integrative review. Psychol Bull 109, 163–203.

Marek, S., and Dosenbach, N.U.F. (2018). The frontoparietal network: function, electrophysiology, and importance of individual precision mapping. Dialogues Clin Neurosci 20, 133–140.

Mayr, U., Awh, E., and Laurey, P. (2003). Conflict adaptation effects in the absence of executive control. Nat Neurosci 6, 450–452.

Menon, V., and Uddin, L.Q. (2010). Saliency, switching, attention and control: a network model of insula function. Brain Struct Funct 214, 655–667.

Milham, M.P., and Banich, M.T. (2005). Anterior cingulate cortex: an fMRI analysis of conflict specificity and functional differentiation. Hum Brain Mapp 25, 328–335.

Miller, E.K., and Cohen, J.D. (2001). An integrative theory of prefrontal cortex function. Annu Rev Neurosci 24, 167–202.

Mitra, P., and Bokil, H. (2008). Observed brain dynamics (New York ; Oxford: Oxford University Press).

Nakamura, K., Roesch, M.R., and Olson, C.R. (2005). Neuronal activity in macaque SEF and ACC during performance of tasks involving conflict. J Neurophysiol 93, 884–908.

Norman, Y., Yeagle, E.M., Khuvis, S., Harel, M., Mehta, A.D., and Malach, R. (2019). Hippocampal sharp-wave ripples linked to visual episodic recollection in humans. Science 365.

Oehrn, C.R., Hanslmayr, S., Fell, J., Deuker, L., Kremers, N.A., Do Lam, A.T., Elger, C.E., and Axmacher, N. (2014). Neural communication patterns underlying conflict detection, resolution, and adaptation. J Neurosci 34, 10438–10452.

Parris, B.A., Wadsley, M.G., Hasshim, N., Benattayallah, A., Augustinova, M., and Ferrand, L. (2019). An fMRI Study of Response and Semantic Conflict in the Stroop Task. Front Psychol 10, 2426.

Reuter, M., Schmansky, N.J., Rosas, H.D., and Fischl, B. (2012). Within-subject template estimation for unbiased longitudinal image analysis. Neuroimage 61, 1402–1418.

Ridderinkhof, K.R., Ullsperger, M., Crone, E.A., and Nieuwenhuis, S. (2004). The role of the medial frontal cortex in cognitive control. Science 306, 443–447.

Rigotti, M., Barak, O., Warden, M.R., Wang, X.J., Daw, N.D., Miller, E.K., and Fusi, S. (2013). The importance of mixed selectivity in complex cognitive tasks. Nature 497, 585–590.

Robertson, J.A., Thomas, A.W., Prato, F.S., Johansson, M., and Nittby, H. (2014). Simultaneous fMRI and EEG during the multi-source interference task. PLoS One 9, e114599.

Sani, I., Stemmann, H., Caron, B., Bullock, D., Stemmler, T., Fahle, M., Pestilli, F., and Freiwald, W.A. (2021). The human endogenous attentional control network includes a ventro-temporal cortical node. Nat Commun 12, 360.

Shenhav, A., Botvinick, M.M., and Cohen, J.D. (2013). The expected value of control: an integrative theory of anterior cingulate cortex function. Neuron 79, 217–240.

Sheth, S.A., Mian, M.K., Patel, S.R., Asaad, W.F., Williams, Z.M., Dougherty, D.D., Bush, G., and Eskandar, E.N. (2012). Human dorsal anterior cingulate cortex neurons mediate ongoing behavioural adaptation. Nature 488, 218–221.

Simon, J.R., and Berbaum, K. (1990). Effect of conflicting cues on information processing: the ‘Stroop effect’ vs. the ‘Simon effect’. Acta Psychol (Amst) 73, 159–170.

Smith, E.H., Horga, G., Yates, M.J., Mikell, C.B., Banks, G.P., Pathak, Y.J., Schevon, C.A., McKhann, G.M., 2nd, Hayden, B.Y., Botvinick, M.M., et al. (2019). Widespread temporal coding of cognitive control in the human prefrontal cortex. Nat Neurosci 22, 1883–1891.

Stroop, J.R. (1935). Studies of interference in serial verbal reactions (Nashville, Tenn.: George Peabody College for Teachers), pp. 19 p.

Tang, H., Yu, H.Y., Chou, C.C., Crone, N.E., Madsen, J.R., Anderson, W.S., and Kreiman, G. (2016). Cascade of neural processing orchestrates cognitive control in human frontal cortex. Elife 5.

Widge, A.S., Heilbronner, S.R., and Hayden, B.Y. (2019). Prefrontal cortex and cognitive control: new insights from human electrophysiology. F1000Res 8.

Wu, J., Ngo, G.H., Greve, D., Li, J., He, T., Fischl, B., Eickhoff, S.B., and Yeo, B.T.T. (2018). Accurate nonlinear mapping between MNI volumetric and FreeSurfer surface coordinate systems. Hum Brain Mapp 39, 3793–3808.

Zilverstand, A., Huang, A.S., Alia-Klein, N., and Goldstein, R.Z. (2018). Neuroimaging Impaired Response Inhibition and Salience Attribution in Human Drug Addiction: A Systematic Review. Neuron 98, 886–903.

